# Identification of a myofibroblast differentiation program during neonatal lung development

**DOI:** 10.1101/2023.12.28.573370

**Authors:** Yongjun Yin, Jeffrey R. Koenitzer, Debabrata Patra, Sabine Dietmann, Peter Bayguinov, Andrew S. Hagan, David M. Ornitz

**Author notes:** Current address, Novartis Biomedical Research, San Diego, 92121 (Andrew S. Hagan). Correspondence: Department of Developmental Biology, 3905 South Bldg. (MSC 8103-0003-03). Co-second authors (Jeffrey R. Koenitzer, Debabrata Patra).

## Abstract

Alveologenesis is the final stage of lung development in which the internal surface area of the lung is increased to facilitate efficient gas exchange in the mature organism. The first phase of alveologenesis involves the formation of septal ridges (secondary septae) and the second phase involves thinning of the alveolar septa. Within secondary septa, mesenchymal cells include a transient population of alveolar myofibroblasts (MyoFB) and a stable but poorly described population of lipid rich cells that have been referred to as lipofibroblasts or matrix fibroblasts (MatFB). Using a unique *Fgf18^CreER^* lineage trace mouse line, cell sorting, single cell RNA sequencing, and primary cell culture, we have identified multiple subtypes of mesenchymal cells in the neonatal lung, including an immature progenitor cell that gives rise to mature MyoFB. We also show that the endogenous and targeted ROSA26 locus serves as a sensitive reporter for MyoFB maturation. These studies identify a myofibroblast differentiation program that is distinct form other mesenchymal cells types and increases the known repertoire of mesenchymal cell types in the neonatal lung.

**Summary Statement:** During primary alveologenesis, alveolar myofibroblasts comprise a distinct proliferative mesenchymal lineage that matures and populates emerging secondary septa.

## Introduction

Alveologenesis is the final stage of lung development where the gas exchange surface area of the lung is increased by subdivision of alveolar saccules followed by thinning of the secondary septae (Rippa et al., 2021). This postnatal developmental process is essential to forming a fully functional adult lung. Impaired alveologenesis is a major cause of morbidity in extremely preterm infants and often results in respiratory distress syndrome which can lead to bronchopulmonary dysplasia (BPD), one of the most significant chronic morbidities affecting nearly half of extreme preterm infants born between 22 and 28 weeks of gestation (Bancalari and Jain, 2019; Shah et al., 2012; Stoll et al., 2015; Whitsett and Weaver, 2015).

Alveologenesis occurs in several phases (Amy et al., 1977; Bostrom et al., 2002; Mund et al., 2008; Rippa et al., 2021; Vila Ellis and Chen, 2021). In the mouse, an initial expansion of alveolar surface area occurs between embryonic day 16.5 (E16.5) and postnatal day 2 (P2) in the absence of alveolar myofibroblasts (MyoFB) and is referred to as the initial or saccular phase. This is followed by the classical or first phase of alveologenesis from P3 to ∼P14, in which MyoFB populate septal ridges, also referred to as secondary crests or secondary septae, that partition the alveolar saccule and thus expand the surface area of the lung. The second phase of classical alveologenesis occurs from ∼P15 to ∼P36 in the absence of MyoFB. During this phase, alveolar size and number continue to increase, and importantly, the septal walls become significantly thinner to facilitate efficient gas exchange.

Septal ridges contain multiple mesenchymal cell types, capillaries, and an elastin rich extracellular matrix (ECM) and are covered by alveolar type 1 (AT1) and alveolar type 2 (AT2) cells (Branchfield et al., 2016; Guo et al., 2023; Yang et al., 2016). Of the mesenchymal cells within the septal ridge, myofibroblasts, also called secondary crest myofibroblasts or alveolar myofibroblasts (MyoFB, SCMF), produce high levels of elastin, and together with their intrinsic contractile properties are essential for secondary septation (Endale et al., 2017; Guo et al., 2023; Li et al., 2019; Li et al., 2015; Lindahl et al., 1997; Narvaez Del Pilar et al., 2022; Sun et al., 2022; Vila Ellis and Chen, 2021; Zepp et al., 2021). The other major mesenchymal cell type is the matrix fibroblast (MatFB) which contains a large subpopulation of lipid-rich cells (lipofibroblasts) and other poorly defined mesenchymal cell types (Riccetti et al., 2020; Rippa et al., 2021). Although understanding the function of the lipofibroblasts is in its early stages (Park et al., 2019), one function is thought to support surfactant and phospholipid synthesis by adjacent AT2 cells (McGowan and Torday, 1997). MatFB also express FGF10, which in the adult lung signals through FGFR2b and is necessary for survival of AT2 cells (Dorry et al., 2020; Liberti et al., 2021; Yuan et al., 2019). It is hypothesized that FGF10 is also necessary to maintain AT2 cells during alveologenesis (Chao et al., 2016; El Agha et al., 2014).

The MyoFB is considered a transient cell population that functions during the first phase of alveologenesis. Genes that are highly expressed in MyoFB, such as ⍺SMA (*Acta2*), are greatly reduced by the end of the first phase of alveologenesis (!P14) suggesting that MyoFB are lost, or alternatively differentiate into another cell type, following their functional role in alveologenesis (Branchfield et al., 2016; McGowan et al., 2008; Riccetti et al., 2020; Yamada et al., 2005). Additionally, several genetic lineage tracing studies have documented that MyoFB are lost by the end of the first phase of alveologenesis and are largely absent from the adult lung (Duong et al., 2022; Gao et al., 2022; Hagan et al., 2020; Li et al., 2015; Narvaez Del Pilar et al., 2022; Zepp et al., 2021).

Several genetic tools have been developed to label in real time and lineage trace mesenchymal cells in the mouse lung (Riccetti et al., 2020). *Pdgfra^EGFP^*(*Pdgfra^tm11(EGFP)Sor/J^*) expresses a nuclear localized H2BEGFP (GFP) gene in a broad range of mesenchymal cells under the control of the endogenous *Pdgfra* locus (Hamilton et al., 2003). In the neonatal lung, the level of GFP fluorescence intensity, as a proxy for *Pdgfra* expression, varies, with low levels (*Pdgfra-*GFP^Low^) observed in MatFB and high levels (*Pdgfra-*GFP^High^) in MyoFB (Endale et al., 2017; Kimani et al., 2009; McGowan and McCoy, 2014; Zepp et al., 2021). Similarly, the adult lung also contains *Pdgfra-*GFP^Low^ and *Pdgfra-*GFP^High^ populations, although the labeled cell types differ from those in the neonatal lung (Green et al., 2016). In the neonatal lung, the different intensities of GFP fluorescence allow sorting dissociated lung cells to isolate MatFB and MyoFB.

Several conditional reporters have also been used to lineage trace lung mesenchymal cells. *Gli1* is expressed in both MyoFB and MatFB at P7 (LungGENS P7 dropseq data) (Du et al., 2015; Du et al., 2017). Activation of a the *Gli1^CreER^; ROSA^mT/mG^* fluorescent genetic reporter mouse at E10.5-12.5 labeled nearly all MyoFB at P14, but also a significant number of PLIN2 positive MatFB, suggesting that at early embryonic time points a common mesenchymal progenitor can give rise to both MyoFB and MatFB (Ahn and Joyner, 2004; Li et al., 2015; Moiseenko et al., 2017; Muzumdar et al., 2007). Activation of *Gli1^CreER^*at P2, P2-5, or P5-6 labeled nearly all MyoFB when examined at P11-15 (Gao et al., 2022; Moiseenko et al., 2017); however, also labeled a large fraction (>20%) of MatFB (at P9) (Hagan et al., 2020) and pericytes, smooth muscle cells, and MatFB (at P14) (Gao et al., 2022). Lineage tracing with *Pdgfra^rtTA^*; TetO-cre; *ROSA^TDTomato^*, activated continuously from E9.5-P7 or from P0-P7, primarily marked MyoFB and a small percentage of MatFB and endothelial cells (Li et al., 2018).

A combination of *Acta2^DsRed^*and *Pdgfra^EGFP^* have also been used to isolate MyoFB (SCMF) and MatFB at P7 (Zepp et al., 2021). Purified cells were used to demonstrate contractile properties of MyoFB, but not MatFB *in vitro*. *Fgf18* is expressed in MyoFB and AT1 cells but not in MatFB (Hagan et al., 2020; McGowan and McCoy, 2015; Ruiz-Camp and Morty, 2015). Consistent with this, activation of FGF18 (TDT) (*Fgf18^CreER^; ROSA^TDTomato^*) from P5-8 labeled nearly all MyoFB and most AT1 cells at P9 with TDTomato (Hagan et al., 2019; Hagan et al., 2020). By P21, most lineage traced MyoFB were no longer detected; however, lineage labeled AT1 cells persisted.

Here, we use a combination of spatial localization, cell sorting, single cell RNA sequencing (scRNA seq), and primary culture of sorted mesenchymal cells to identify a spectrum of mesenchymal cell types in the neonatal lung. This study shows a separation of MyoFB and MatFB lineages in the neonatal lung and identifies a progression from immature progenitors to mature MyoFB in sequence data, immunostaining patterns, and in *in vitro* cell culture. We also uncovered an unanticipated regulation of the endogenous and targeted *ROSA26* locus (*Gt(ROSA)26Sor*) as a sensitive reporter for MyoFB maturation. These studies increase the known repertoire of mesenchymal cell types in the neonatal lung and provide a neonatal time point for comparison with embryonic and adult lung mesenchymal cell subtypes with implications for activation of developmental programs that could function during lung injury and repair.

## Results

### Localization of myofibroblasts and matrix fibroblasts relative to the elastin lung scaffold in the neonatal mouse

*Fgf18^CreERT2^; ROSA^TDTomato^*; *Pdgfra^EGFP^*mice can be used to label MyoFB, AT1 cells and mesothelial cells in red (*Fgf18*-TDTomato lineage), and MyoFB and MatFB in green in the neonatal mouse lung (Endale et al., 2017; Hagan et al., 2020; Li et al., 2018; McGowan et al., 2008). *Pdgfra^EGFP^* labels MatFB with lower GFP intensity and MyoFB with higher GFP intensity (Kimani et al., 2009; McGowan and McCoy, 2014). To localize these cells relative to the P7 neonatal elastin lung scaffold, we used CLARITY to optically clear lungs in combination with a chemical stain (Alexa Fluor 633 Hydrazide) that selectively stains elastin to visualize alveolar entry rings and septal ridges (Fig. 1A-D) (Shen et al., 2012). Confocal imaging of Alexa Fluor 633 Hydrazide-stained lungs showed a dense elastin matrix in the mesothelial surface (top 5 µm) and elastin bundles outlining alveolar entry rings in the underlying tissue. *Pdgfra^EGFP^* (GFP) labeled the nucleus of cells that were distributed throughout the lung, but excluded from the mesothelium (Fig. 1E, F). Induction of the FGF18 (TDT) (*Fgf18^CreERT2^; ROSA^TDTomato^*; *Pdgfra^EGFP^*) lineage trace with tamoxifen from P2-P5 showed abundant TDTomato-labeled cells throughout the lung and in the mesothelium (Fig. 1A, E-G). Merged color channels clearly identify at least 4 distinct cell types: MatFB (green, PDGFRA(GFP)^Low^), MyoFB (green, PDGFRA(GFP)^High^ and yellow, PDGFRA(GFP)^High^ + FGF18 (TDT)), AT1 cells and mesothelial cells (red, FGF18 (TDT)). High magnification images show MyoFB in close proximity to the elastin bundles and immature FGF18 (TDT)-negative MyoFB (see below) and MatFB towards the base of septal ridges (Fig. 1g, h, Supplemental Movie 1). This pattern of cellular localization is consistent with numerous studies using a variety of cellular markers that label MyoFB and MatFB (Branchfield et al., 2016; Endale et al., 2017; Hagan et al., 2020; Kimani et al., 2009; McGowan et al., 2008; McGowan and McCoy, 2014; Riccetti et al., 2020).

**Figure 1.**
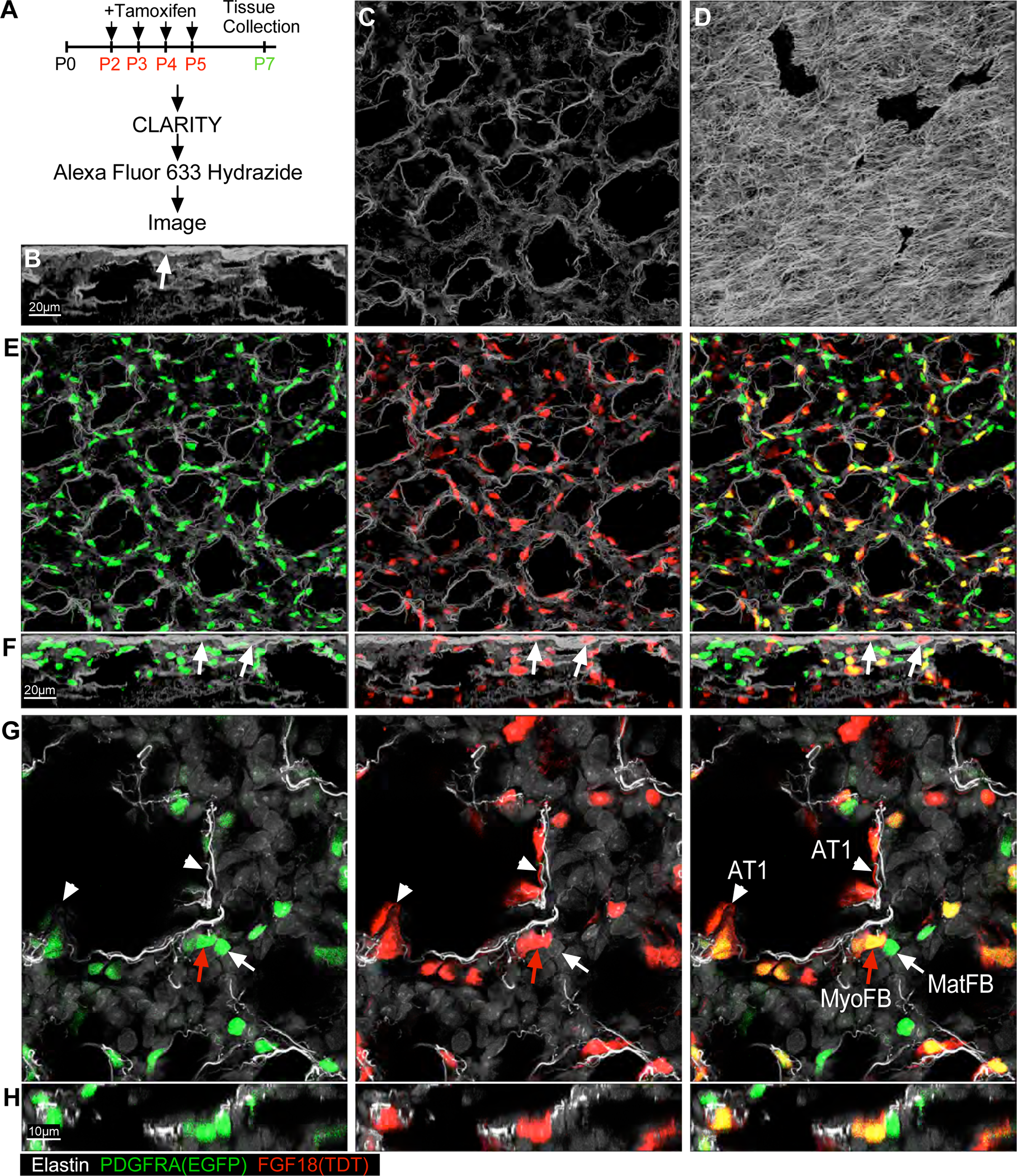
High resolution imaging of elastin and lineage labeled cells in the postnatal day 7 mouse lung. **A**. Experimental plan for imaging *Fgf18^CreER^; ROSA^TDTomato^*; *Pdgfra^EGFP^* cleared lung tissue using CLARITY. **B-H**. XY and Z plane maximum intensity projection (MIP) images of Elastin (Alexa Fluor 633 Hydrazide-stained lung, white), *Fgf18^CreER^; ROSA^TDTomato^* (Red), and *Pdgfra^EGFP^* (Green) fluorescent proteins. **B**. Z plane image showing a dense elastin matrix on the lung surface (arrow). **C**. XY plane image showing the elastin matrix, 18-38 µm below the lung surface. **D**. XY plane image showing the elastin matrix in the mesothelium (top 0-5 µm). **E, F.** Low magnification images showing XY (**E**) and Z (**F**) plane images of *Fgf18* lineage (Red) and *Pdgfra^EGFP^* expression (Green). Arrows in F show *Fgf18* lineage labeled cells in the mesothelium. **G**, **H**. High magnification images showing the spatial relationship of labeled cells. AT1 cells (white arrowhead), MyoFB (red arrow), MatFB (white arrow). Representative images from at least three mice are shown.

### Identification of mesenchymal cell types in the neonatal lung

To identify the repertoire of mesenchymal cells in the neonatal lung, *Fgf18^CreERT2/+^; ROSA^TDTomato/+^*; *Pdgfra^EGFP/+^*mice were induced with tamoxifen from P2-P5, and lungs were dissociated for cell sorting on P7 (Fig. 1A, 2A). FACS identified four groups of cells (T, C1, C2, C3) (Fig. 2A). Group T (TDTomato^+^, GFP^−^) is enriched in AT1 cells and mesothelial cells as shown by increased relative levels of podoplanin (*Pdpn*) and Wilms tumor 1 homolog (*Wt1*) expression, respectively (supplemental Fig. 1). Sorted C1 (TDTomato^−^, GFP^low^), C2 (TDTomato^−^, GFP^High^) and C3 (TDTomato^+^, GFP^High^) cells were individually subjected to scRNA seq using the 10x Genomics platform (Fig. 2A, B). After filtering data, preliminary clustering, and removal of a small number of contaminating hematopoietic and AT2 cells (109 from *Fgf18^CreERT2/+^* samples), we analyzed 1284, 3347, 1511 cells from samples C1, C2, and C3, respectively (Supplemental Table 1A). The sequence data, pooled along with a similar data set from *Fgf18* conditional knockout mice (*Fgf18^CreERT2/f^*), was used for cluster analysis and cell type identification. All data presented here is from lineage traced lungs that are heterozygous for *Fgf18* (*Fgf18^CreERT2/+^*).

**Figure 2.**
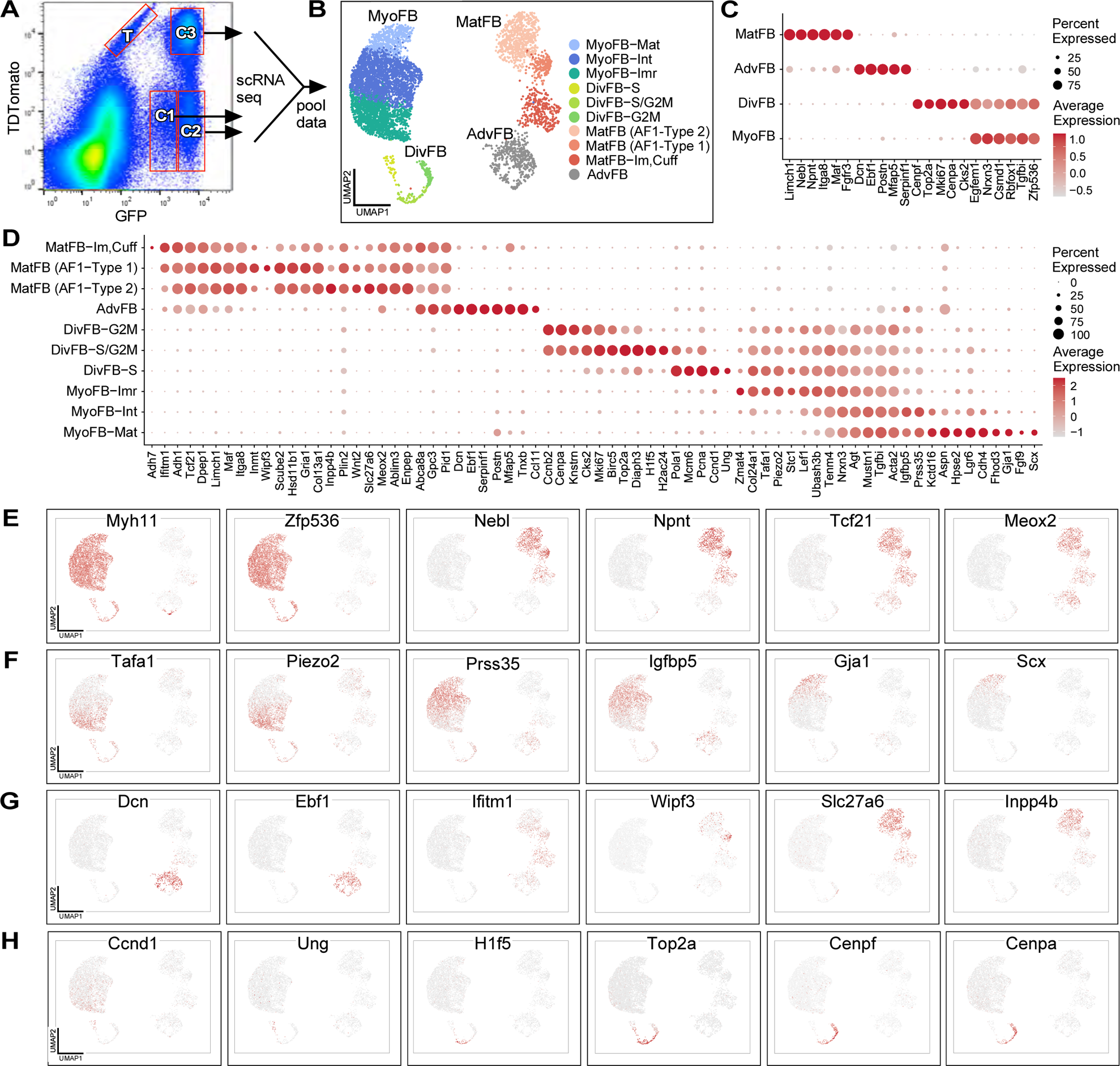
Identification of mesenchymal cell populations in the postnatal day 7 mouse lung. **A**. Whole lung from P7 *Fgf18^CreER^; ROSA^TDTomato^*; *Pdgfra^EGFP^* mice (treated with tamoxifen from P2-P5) were dissociated and sorted for GFP and TDTomato. Four cell types were collected: T (TDTomato^+^, GFP^−^); C1 (TDTomato^−^, GFP^Low^), C2 (TDTomato^−^, GFP^High^) and C3 (TDTomato^+^, GFP^High^). C1, C2, and C3 cells were subjected to scRNA seq. **B**. Cell clustering identified four primary cell subtypes (MyoFB, MatFB, DivFB and AdvFB) that could be further subclustered as indicated. **C**. Dotplot showing marker genes highly selective for expression in the four primary cell types. **D**. Dotplot showing marker genes selective for expression in the 10 subclusters. **E**. UMAP showing examples of genes that are selective for MyoFB and MatFB. **F**. UMAP showing examples of genes selective for subclusters of MyoFB. **G**. UMAP showing examples of genes selective for AdvFB and subclusters of MatFB. **H**. UMAP showing examples of genes selective for subclusters of DivFB.

The identified cell clusters clearly segregated into MyoFB, MatFB, adventitial (AdvFB) and proliferating (DivFB) cell clusters (Fig. 2B, Table 1). These clusters could be distinguished by markers such as *Myh11* and *Zfp536* which were highly specific for MyoFB, *Npnt* and *Nebl* which were highly specific for MatFB, and *Dcn* and *Ebf1* which were highly specific for AdvFB (Fig. 2C-G). *Tcf21* and *Meox2* mark MatFB but are also present in AdvFB (Fig. 2C-E). *Plin2* (*Adrp*), a marker of lipofibroblasts, is expressed at low levels in most cells, but enriched in MatFB (Fig. 2d, Table 1, Supplemental Table 1B).

**Table 1.**
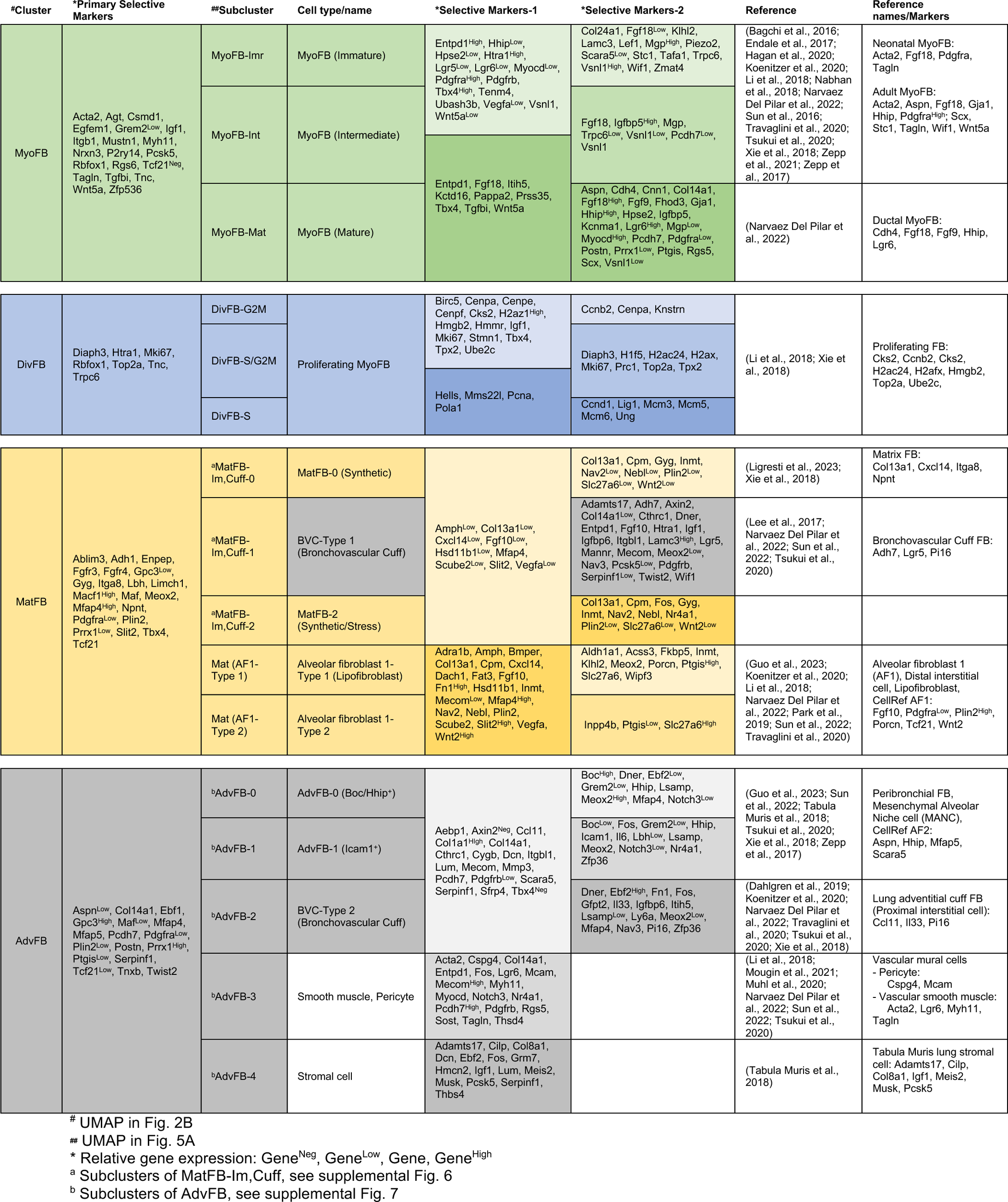
Selective gene expression markers for postnatal day 7 lung mesenchymal fibroblasts.

Clusters enriched for proliferating cells (DivFB) were also identified and the majority of these cells shared markers with MyoFB (Fig. 2C, Table 1). Proliferating cells could be separated into S phase, mixed S/G2M, and G2M phase clusters using cell cycle scoring (Seurat) (Fig. 2B-D, H, Supplemental Fig. 2, Supplemental Table 1B) (Tirosh et al., 2016). Subclusters of proliferating cells could be distinguished by markers such as *Ung*, *H1f5*, *Cenpa,* and *Top2a* (Fig. 2C, D, H). Analysis of proliferating cells grouped by the original cell sorting sample showed that C2 cells (TDTomato^−^, GFP^High^) were enriched for DivFB compared to C1 and C3 samples (Supplemental Fig. 2).

### Identification of myofibroblast subtypes

Within the MyoFB group (*Myh11+, Tagln+*), we could identify three types of cells which we designated as immature (MyoFB-Imr), intermediate (MyoFB-Int), and mature (MyoFB-Mat) myofibroblasts (Fig. 2B-D, F, Table 1, Supplemental Table 1B). MyoFB-Imr cells were identified by high expression of *Zmat4, Tafa1, Lef1, Ubash3b,* and *Piezo2,* and high levels of *Pdgfra* and *Pdgfra^EGFP^* (Fig. 2D, F, Supplemental Fig. 3). MyoFB-Mat cells were enriched for *Aspn*, *Cdh4, Col14a1, Fgf9, Fgf18, Gja1* (Cx43)*, Hhip,* and *Lgr6* (Fig. 2D, F, Supplemental Fig. 3). MyoFB-Mat cells show similar gene expression with *Cdh4/Hhip/Lgr6* ductal myofibroblasts and MyoFB-Imr cells appear similar to PDGFRA-high alveolar myofibroblasts identified by Narvaez Del Pilar et al. (Narvaez Del Pilar et al., 2022) (Table 1, Supplemental Fig. 3a). Additionally, we find that a subset of the MyoFB-Mat cells expressed high levels of *Scx* and *Fgf9* (Fig. 2D, F, Supplemental Fig. 3B). We also identified an intermediate cell population (MyoFB-Int) characterized by higher levels of *Prss35* and *Igfbp5* (Fig. 2D, F). We posit that these cells represent a transitional state between immature and mature MyoFB. *Prss35* is interesting as it is a profibrotic matrisomal protein that regulates cell proliferation in response to hyperosmotic stress (Sanger et al., 2023).

Immunohistochemical staining was used to analyze the location of markers that identify immature and mature MyoFB cell types (Fig. 3, Supplemental Fig. 4). Mature MyoFB within septal ridges could be identified that co-express FGF18 (TDT), ⍺SMA, and Cx43 (*Gja1*) (Fig. 3A). FGF9 expression was analyzed in the lung of *Fgf9^βGal^*mice (Huh et al., 2015) using an antibody to βGalactosidase. This showed co-localization of FGF9 (βGal), Cx43 and ⍺SMA in MyoFB-Mat cells within septal ridges (Fig. 3B). Scx lineage cells, identified by immunostaining SCX (TDT) mice (Scx-Cre; *ROSA^TDTomato^*) (Sugimoto et al., 2013) for TDTomato, showed co-localization with ⍺SMA and Cx43 in MyoFB-Mat cells (Fig. 3C). CD34 is a transmembrane glycoprotein that is expressed in mesenchymal stem cells, muscle-derived stem cells, hematopoietic progenitor cells and stem/progenitor cells of other tissues (Bose and Shenoy, 2016; Diaz-Flores et al., 2014; He et al., 2013; Sidney et al., 2014). In MyoFB, *Cd34* is expressed at high levels in MyoFB-Imr and low levels in MyoFB-Mat cells (Supplemental Fig. 3B). Consistent with expression in progenitor cells, CD34^+^, Cx43^+^, TAGLN^+^ immature MyoFB and CD34^−^, Cx43^+^, TAGLN^+^ mature MyoFBs were both identified in the developing alveoli (Fig. 3D). Similarly, FGF18 (TDT)^−^, Cx43^+^, CD34^+^ MyoFB-Imr cells were distinct from FGF18 (TDT)^+^, Cx43^+^, CD34^−^ MyoFB-Mat cells (Fig. 3E). *Lef1* is another marker preferentially expressed in MyoFB-Imr cells (Fig. 2D, Supplemental Fig. 3B). Co-immunostaining for FGF9 (βGal) and LEF1 identified LEF1^+^, FGF9 (βGal)^−^ MyoFB-Imr cells, LEF1^+^, FGF9 (βGal)^+^ MyoFB-Int cells and LEF1^−^, FGF9 (βGal)^+^ MyoFB-Mat cells (Supplemental Fig. 4A). Co-immunostaining with Cx43 identified Cx43^+^, LEF1^+^ MyoFB-Int cells and Cx43^−^, LEF1^+^ MyoFB-Imr cells (Supplemental Fig. 4B). Consistent with increased proliferation of MyoFB-Imr cells, SCX (TDT)^−^ cells co-immunostained for LEF1 and MKI67, whereas septal tips contained SCX (TDT)^+^, MKI67^−^, LEF1^−^, MyoFB-Mat cells (Supplemental Fig. 4C).

**Figure 3.**
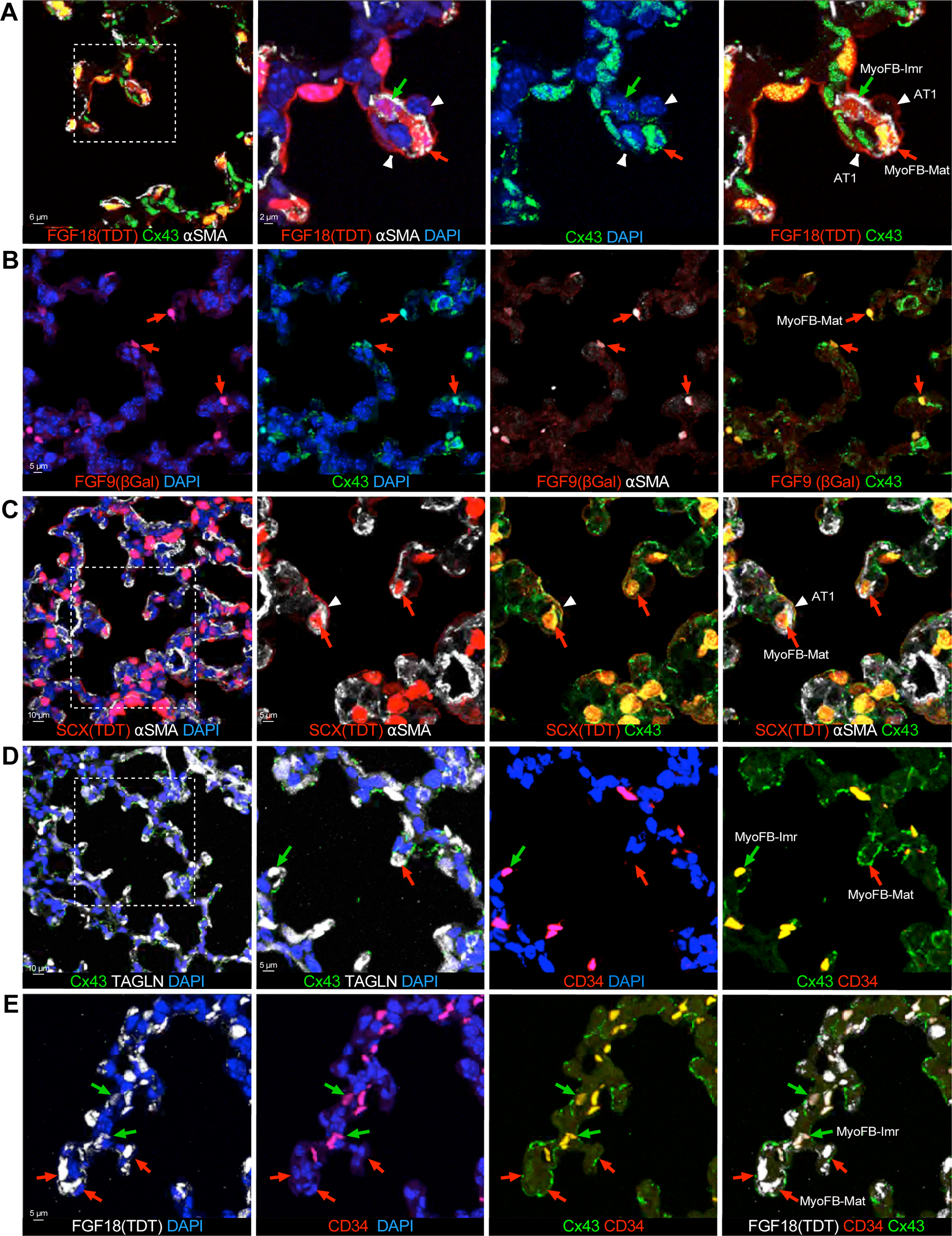
Identification of immature and mature myofibroblast subtypes. **A**. MyoFB-Mat cells (red arrow) within the lung septal ridges are identified by co-staining for Cx43+, ⍺SMA^+^ and FGF18 (TDT)^+^ cells; MyoFB-Imr cells (green arrow) only co-stain for ⍺SMA^+^ and FGF18 (TDT)^+^ but are negative for Cx43 expression. AT1 cells (white arrow heads) are FGF18 (TDT)^+^ and have AT1 cell morphology. Dashed box in the left image indicates area shown at higher magnification in the following images. **B**. MyoFB-Mat cells (red arrows) identified in *Fgf9^βGal^* mice show co-localization of FGF9 (βGal), Cx43 and ⍺SMA. **C**. MyoFB-Mat cells (red arrows) identified in SCX (TDT) mice show co-localization of TDT, Cx43, and ⍺SMA. AT1 cells (white arrowhead) also contain SCX (TDT) and Cx43. **D**. TAGLN (SM22) labels all MyoFB. MyoFB-Mat cells (red arrow) co-stain with Cx43 and do not contain CD34. MyoFB-Imr (green arrow) are CD34^+^ and Cx43^+^. In Myo-Imr cells, Cx43 expression is cytosolic and not localized in the cell membrane as in MyoFB-Mat cells. **E**. MyoFB-Mat cells (red arrows) contain FGF18 (TDT) and Cx43, but do not stain for CD34. MyoFB-Imr cells (green arrows) are Cx43^+^, CD34^+^, but contain low levels of FGF18 (TDT). Representative images from at least three mice are shown.

RNAseq data shows *Pdgfra^EGFP^* (GFP) at higher levels in MyoFB-Imr cells and lower levels in MyoFB-Mat cells (Supplemental Fig. 3A). The endogenous *Pdgfra* and *Pdgfrb* transcripts have a similar profile. However, *Fgf18* has the opposite pattern, with low levels of FGF18 (TDT) in MyoFB-Imr and higher levels in MyoFB-Mat cells (Supplemental Fig. 3B). Co-immunostaining for the *Fgf18* lineage, FGF18 (TDT), GFP, and Cx43 identified FGF18 (TDT)^+^, GFP^High^, Cx43^+^ MyoFB-Mat cells, and FGF18 (TDT)^+^, GFP^Low^, Cx43^+^ MyoFB-Int cells (Supplemental Fig. 4D). To confirm the expression pattern of *Pdgfra^EGFP^* (GFP), we compared its expression with endogenous PDGFRA and ⍺SMA (*Acta2*). Co-immunostaining showed co-localization of both markers in compact MyoFB-Imr cells and in larger MyoFB-Mat cells (Supplemental Fig. 4E).

To identify potential functions of MyoFB, differentially expressed genes (DEG) that were enriched in MyoFB-Imr and MyoFB-Mat clusters as well as FGF18 (TDT) negative and positive cells were subjected to GO classification. MyoFB-Imr cells were associated with GO terms that suggest transition to the G1 phase of the cell cycle and GO terms such as cytoplasmic translation and macromolecule biosynthetic process that indicates enhanced biosynthetic capacity, consistent with production of extracellular matrix (ECM) proteins (Supplemental Fig. 5A). The GO terms associated with MyoFB-Imr cells were most similar to FGF18 (TDT)^−^ MyoFB (Supplemental Fig. 5B). By contrast, MyoFB-Mat cells expressed genes involved in organization of the ECM and, interestingly, are a source of secreted signaling molecules that might regulate cells in the local environment including the potential regulation and patterning of angiogenesis (Supplemental Fig. 5C). The GO terms associated with MyoFB-Mat cells were most similar to FGF18 (TDT)^+^ MyoFB (Supplemental Fig. 5D). These data suggest that FGF18 (TDT) expression is an indicator of MyoFB maturation.

### Identification of matrix fibroblast subtypes

Within the MatFB group, we could identify three large distinct clusters (Fig. 2B-D, Table 1, Supplemental Table 1B). MatFB-Im,Cuff cells include an immature matrix fibroblast and a population of bronchovascular cuff (BVC) adventitial fibroblasts (detailed below). This cluster of cells was identified by higher expression of *Ifitm1* and *Adh1* (Fig. 2B, D, G, Table 1). The other two MatFB clusters were most similar to previously defined alveolar fibroblast 1 cells (AF1) in adult mice which are marked by expression of *Tcf21* and *Wnt2* (Sun et al., 2022) and more specifically by *Bmper, Col13a1, Fat3*, and *Fgfr4* (Koenitzer et al., 2020). We designated these AF1 cells as MatFB (AF1-Type 1) and MatFB (AF1-Type 2). MatFB (AF1-Type 1) cells express high levels of *Wipf3* and *Scube2* and MatFB (AF1-Type 2) cells express high levels of *Wnt2*, *Slc27a6*, and *Inpp4b* (Fig. 2B, D, G, Table 1).

The MatFB-Im,Cuff cluster is heterogeneous, as a subset of these cells specifically express *Adh7* (Fig. 2D, Supplemental Fig. 6A, Table 1, Supplemental Table 1B), a gene found previously to mark a unique population of cytokine-deficient adventitial fibroblasts (Tsukui et al., 2020). These cells also express *Dner* and *Twist2*, but do not express *Dcn*, *Ebf1, Ccl11,* or *Il33*, genes expressed in classical adventitial fibroblasts (Fig. 2C, D, G, Table 1, Supplemental Figs. 6A, 7A) (Dahlgren et al., 2019; Narvaez Del Pilar et al., 2022; Tsukui et al., 2020). Subclustering of MatFB-Im,Cuff cells identified three subclusters (Table 1, Supplemental Fig. 6B, Supplemental Table 1C). Subcluster 0 was distinguished from subclusters 1 and 2 by expression of high levels of *Nebl* and *Nav2* (regulates cell migration) (Schmidt et al., 2009) and enrichment for ribosomal and ECM genes. These cells appear distinct from NEBL+ pathological MyoFB seen in IPF lungs which also express high levels of *MYO10* and *RYR2* (Adams et al., 2020). Subcluster 0 cells were designated as immature synthetic cells, MatFB-0 (Synthetic) (see gene ontology below, Supplemental Fig. 5E).

Subcluster 1 cells expresses *Adh7*, *Col14a1, Dner, Pdgfrb,* and *Twist2,* and do not express Il33. These cells are most similar to adventitial or BVC fibroblasts identified in P7 and adult lung (Narvaez Del Pilar et al., 2022; Tsukui et al., 2020). We designate these as BVC-Type 1 fibroblasts (Table 1, Supplemental Fig. 6, Supplemental Table 1C). Subcluster 2 appears to contain stress response factors (e.g., *Fos, Nr4a1, Atf3*) and is likely not a unique fibroblast population. The location of MatFB-Im,Cuff subclusters relative to all other clusters is shown in Fig. 5A.

Immunohistochemical staining was used to assess the location and co-localization of MEOX2, a transcription factor expressed in MatFB and AdvFB, but not in MyoFB (Figs. 2D, 4, Table 1) (Narvaez Del Pilar et al., 2022). In the distal lung, MEOX2 was expressed in cells that do not express FGF18 (TDT) (Fig. 4A-C). Consistent with previous studies, MEOX2 co-localized with *Pdgfra-*GFP^Low^ expressing MatFB but not with *Pdgfra-*GFP and ⍺SMA expressing MyoFB (Fig. 4A, B). In the more proximal lung, MEOX2 was expressed in the bronchovascular cuff region, and was not expressed in FGF18 (TDT) and ⍺SMA expressing airway and vascular smooth muscle cells (Fig. 4C, D).

**Figure 4.**
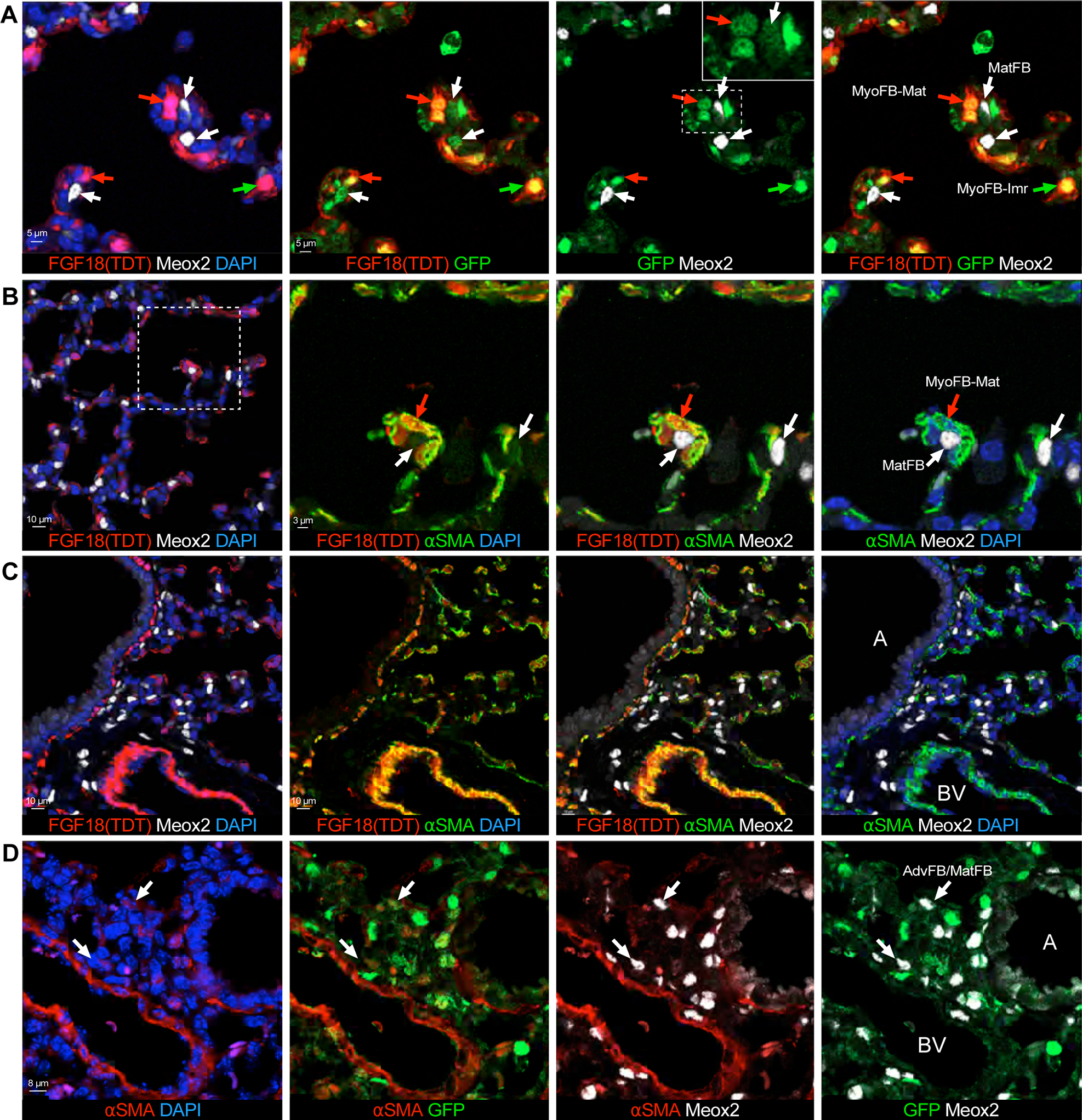
Identification of adventitial and matrix fibroblasts. **A**. MatFB within the lung septal ridges contain low levels of *Pdgfra^EGFP^* (GFP) and MEOX2 (white arrow) and show no overlap with FGF18 (TDT). MyoFB-Mat (red arrow) cells contains FGF18 (TDT) and low levels of *Pdgfra^EGFP^* (GFP), but do not contain MEOX2. MyoFB-Imr cells (green arrow) contain FGF18 (TDT) and high levels of *Pdgfra^EGFP^* (GFP), but do not contain MEOX2. The inset shows a magnification of the dashed box with only the green color channel to demonstrate dim GFP expression in a MEOX2^+^ cell (white arrow). **B**. Dashed box in the left image indicates area shown at higher magnification in the following images. MEOX2 (white arrow) in MatFB do not co-localize with ⍺SMA or FGF18 (TDT) in MyoFB-Mat cells (red arrow). **C**, **D**. Analysis of the bronchovascular cuff region in the proximal lung. **C**. MEOX2^+^ cells are distributed around the airways and vessels and do not co-localize with FGF18 (TDT) and ⍺SMA in vascular and airway smooth muscle. **D**. Adventitial or matrix fibroblasts (AdvFB/MatFB) contain MEOX2 and low levels of *Pdgfra^EGFP^* (GFP) (white arrow) and are present in tissue surrounding the airways and vessels. Representative images from at least three mice are shown.

*<u>GO analysis.</u>* MatFB-Im,Cuff subcluster 1 (BVC-Type 1) cells and AdvFB both express *Dner* and *Twist2* suggesting that these cell types are related (Supplemental Fig. 6A, Table 1). For gene ontogeny analysis of MatFB, we therefore excluded MatFB-Im,Cuff subcluster 1 (BVC-Type 1) cells. MatFB DEGs were determined from a subset of cells that include MatFB-Im,Cuff subcluster 0, MatFB (AF1-Type 1) and MatFB (AF1-Type 2). MatFB-Im,Cuff subcluster 0 cells were associated with GO terms that include cytoplasmic translation, gene expression, and cellular respiration, characteristics of an immature matrix producing cell. We refer to these cells as MatFB-0 (Synthetic) cells (Supplemental Fig. 5E, 6B). MatFB (AF1-Type 1) cells are associated with GO terms that include positive regulation of kinase activity, small GTPase signal transduction, transmembrane RTK signaling pathways, and regulation of endothelial cell migration, suggesting that these cells integrate multiple environmental signals and function to organize the local tissue environment (Supplemental Fig. 5E). MatFB (AF1-Type 2) cells are associated with GO terms that include cell substrate junction, focal adhesion and cellular response to growth factor stimulus, suggesting that these cells interact with adjacent cells through adhesion molecules/junctional proteins and integrate intercellular signals within tissues (Supplemental Fig. 5E).

### Identification of adventitial fibroblast subtypes

The AdvFB group expresses high levels of *Dcn*, *Col14a1,* and *Cthrc1* (Fig. 2D, G, Table 1, Supplemental Fig. 7, Supplemental Table 1B, D). Within this group, five subclusters could be identified (Supplemental Table 1D). *Cthrc1* is expressed at highest levels in clusters 0, 1 and 2, suggesting that these cells may be a reservoir of cells that could contribute to fibrotic lung disease in the adult lung (Tsukui et al., 2020). These subclusters also express *Ccl11* and *Il33* which have been found in cells within the bronchovascular cuffs in adult lung (Fig. 2D, Table 1, Supplemental Fig. 7A, Supplemental Table 1D) (Dahlgren et al., 2019; Tsukui et al., 2020). Subcluster 0 was enriched for expression of *Boc, Hhip,* and *Lsamp,* and subcluster 1 was enriched for expression of *Zfp36* and *Icam1.* We refer to these clusters as Boc/Hhip^+^ and Icam1^+^ AdvFB, respectively. Subcluster 2 was enriched for expression of *Ebf2, Il33,* and *Pi16,* and corresponds to a subtype of adult lung bronchovascular cuff fibroblasts (Tsukui et al., 2020). We designate these as BVC-Type 2 fibroblasts (Table 1, Supplemental Fig. 7, Supplemental Table 1D).

Subcluster 3 (vascular smooth muscle, pericyte) expressed *Acta2, Myh11,* and *Tagln,* markers of smooth muscle, but also contained a subpopulation of cells expressing *Sost* and *Cbr2* that are similar to heart vascular mural cells (Muhl et al., 2020), *Rgs5* and *Cspg4* that mark pericytes (Tsukui et al., 2020), and *Thsd4* (encoding ADAMTSL6), an ECM protein associated with elastic tissues (Mougin et al., 2021). Subcluster 4 cells expressed lung stromal markers, *Col8a1, Igf1, Meis2*, and *Pcsk5*, identified in Tabula Muris, but also specifically expressed *Grm7* and *Thbs4*, not found in the lung in Tabula Muris (Tabula Muris et al., 2018), suggesting a potentially unique lung adventitial cell type.

*<u>GO analysis.</u>* AdvFB DEGs were determined from a subset of cells that include AdvFB subclusters 0, 1, 2 and BVC-Type 1 cells derived from MatFB-Im,Cuff subcluster 1 (BVC-Type 1) (Supplemental Fig. 5F). All of these cells were associated with GO terms that include collagen containing extracellular matrix. AdvFB subcluster 0 (*Boc/Hhip^+^*) cells were most associated with GO terms that include organization of the extracellular environment, cell migration and cell adhesion. AdvFB subcluster 1 (*Icam1^+^*) cells were associated with GO terms that include focal adhesion, and regulation of apoptotic process, proliferation, and migration. AdvFB subcluster 2 (BVC-Type 2) cells may additionally regulate the extracellular environment, including angiogenesis. MatFB-Im,Cuff subcluster 1 (BVC-Type 1) cells have GO terms associated with cell adhesion, extracellular matrix organization, and regulation of cell proliferation.

### Trajectory analysis predicts differentiation of distinct neonatal myofibroblast and matrix fibroblast progenitors

To evaluate potential lineage relationships within neonatal fibroblast populations, we employed multiple trajectory inference approaches. For RNA velocity (La Manno et al., 2018), ratios of spliced to unspliced transcripts for each gene are used to extrapolate temporal relationships among cells. In myofibroblasts, the inferred trajectory supported a differentiation pathway from dividing to immature and ultimately to mature myofibroblasts as projected in UMAP space (Fig. 5A-C). Non-linear dimensional reduction by diffusion components results in diffusion maps (DM) that often recapitulate biological differentiation pathways in scRNA seq data (Haghverdi et al., 2015). Diffusion mapping on a subset of cells including all myofibroblast populations also suggested a linear trajectory from MyoFB-Imr to MyoFB-Mat, further supported by projection of RNA velocity on DM coordinates. Finally, pseudotime analysis was performed using *Slingshot* (Street et al., 2018), with DivFB defined as the starting cluster. This approach also identified MyoFB-Mat as a differentiation endpoint.

**Figure 5.**
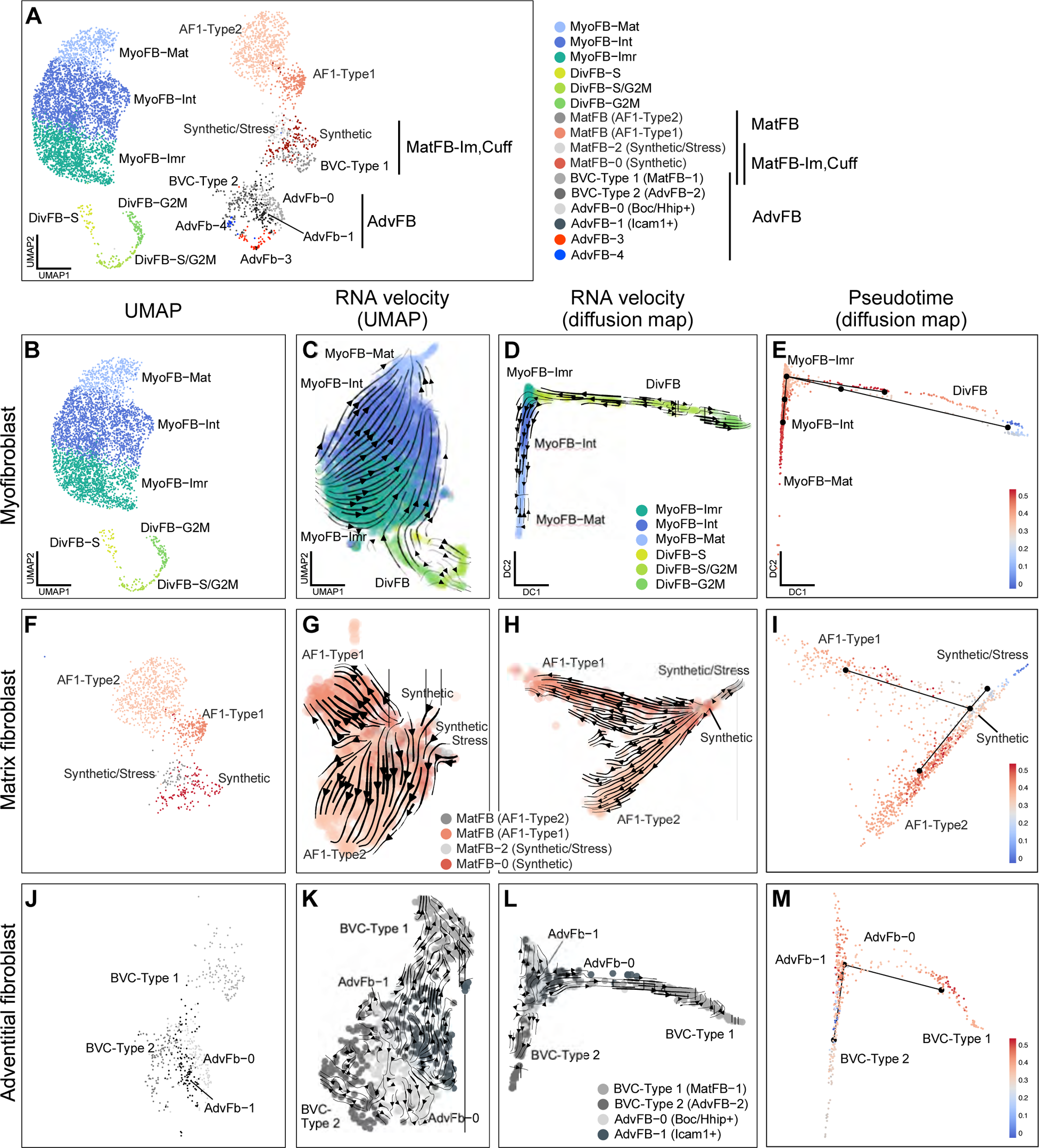
Potential differentiation trajectories for lung mesenchymal cell populations determined by RNA velocity and pseudotime analysis. **A**. UMAP showing a projection of subclusters of MatFB-Im,Cuff cells and AdvFB. **B-E**. Trajectory analysis of P7 myofibroblasts (MyoFB). **B**. UMAP projection of myofibroblast populations and dividing cells. **C**. RNA velocity determined by *scVelo*, indicated as arrow streams on the MyoFB UMAP projection indicating a trajectory from MyoFB-Imr to MyoFB-Mat cells, with arrowheads denoting direction of differentiation. **D**. Diffusion map for MyoFB using the first two diffusion components with overlaid RNA velocity streams. **E**. *Slingshot* calculated pseudotime projected on the same diffusion map with DivFB designated as the starting cluster. **F-I**. Trajectory analysis of MatFB subtypes excluding the BVC-Type 1 subcluster. **F**. UMAP projection of MatFB populations and putative precursors. **G**. RNA velocity stream projected in UMAP space from *scVelo*. **H**. RNA velocity projections on the MatFB diffusion map. **I**. *Slingshot-*calculated pseudotime projected onto the MatFB diffusion map with the starting cluster designated as MatFB-0 (Synthetic). **J-M**. Trajectory analysis of adventitial (AdvFB) and bronchovascular cuff type 1 (BVC-Type 1) fibroblasts. **J**. UMAP projection of the relevant adventitial/cuff clusters. **K, L**. RNA velocity depicted as arrow streams as determined by *scVelo* projected onto the UMAP (**k**) and diffusion map (**L**) coordinates. **M**. *Slingshot-*calculated pseudotime projected onto the diffusion map with AdvFB-0 designated as the starting cluster.

For matrix fibroblasts, a similar analysis was performed on the AF1-Type 1, AF1-Type 2, Synthetic and Synthetic/Stress populations. Proposed trajectories by all methods showed progression from Synthetic cells with AF1-Type 1 and AF1-Type 2 as separate endpoints (Fig. 5B, F-I). By diffusion map, Synthetic/Stress cells largely form a third trajectory away from the Synthetic cluster (Fig. 5H, I), suggesting that these cells are not an intermediate in the development of mature alveolar fibroblast populations. By our classification, adventitial fibroblasts included BVC-Type 1 (MatFB-1) and BVC-Type 2 (AdvFB-2) cells along with AdvFB-0 and AdvFB-1 subclusters. AdvFB-3 and AdvFB-4 were considered unlikely to share a lineage relationship with these subclusters and were thus excluded from this analysis. Trajectory analyses of these cell types as a subset suggest that AdvFB-1 cells are a point of origin that could give rise to the other subtypes, and that BVC-Type 1 and BVC-Type 2 cells represent distinct lineages (Fig. 5L-M). No differentiation pathway linking MyoFB and MatFB was noted, suggesting that at this stage of development myofibroblasts and matrix fibroblasts comprise distinct lineages.

### Intercellular communication

To evaluate intercellular signaling among the full complement of cell types in the developing alveolus including our identified fibroblast subpopulations, we merged and integrated our annotated expression matrix with an existing lung scRNA seq dataset from P7 mouse lung (Fig. 6A) (Zepp et al., 2021) and performed ligand-receptor analysis with CellChat. MyoFB are recipients of an array of extracellular signals including SHH, PDGF, TGFβ, WNT, and NOTCH (Fig. 6B-F), and also act as a source of secreted signaling molecules including FGF18 and WNT5a (Fig. 6F, G). Differential signaling was noted among subpopulations of MatFB and MyoFB as well; among MyoFB, FGF9 production is limited to MyoFB-Mat (Fig. 6G). Similarly, MyoFB-Imr cells are predominantly recipients of FGF18 signaling via FGFR1, while MyoFB-Int and MyoFB-Mat cells are primarily producers of FGF18 (Fig. 6G). Among matrix fibroblasts, few interactions distinguished AF1-Type 1 and AF1-Type 2 cells, though they may differ in their capacity to signal via TGFB2 (Fig. 6C).

**Figure 6.**
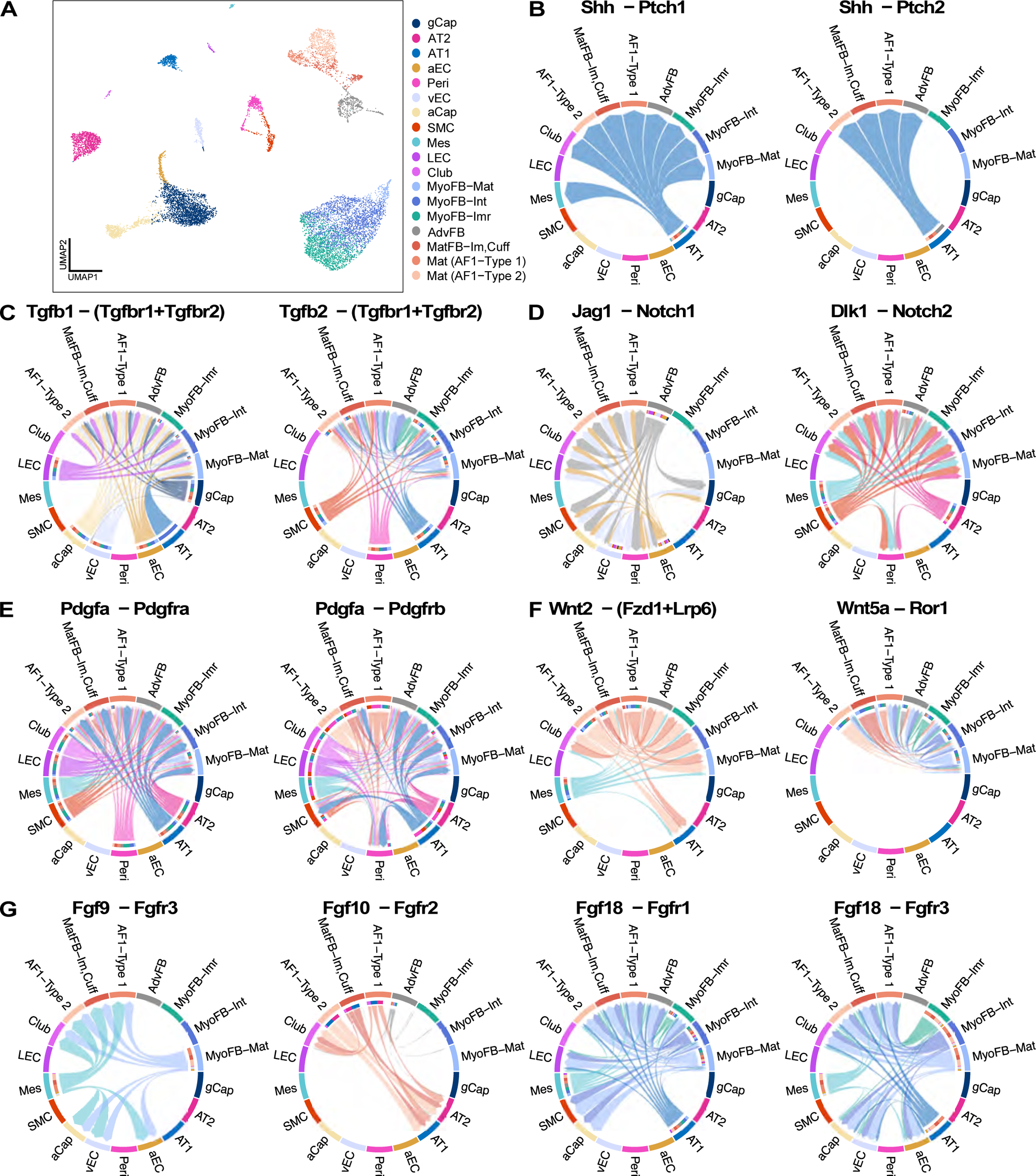
Selected potential intercellular communication pathways between lung mesenchymal cells and other lung cell types identified using cell chat. **A**. UMAP projection of combined Zepp et al. dataset and sorted mesenchymal cells after merge and reclustering, as performed in preparation for CellChat analysis. **B-G**. CellChat chord diagrams for selected signaling pathways. Terminal arrow width corresponds to relative signal strength. **B**. Sonic hedgehog via Patched1 and Patched2, predominantly from AT1 to cells to fibroblast populations. **C**. TGFβ 1 and 2 signaling. **D**. Notch signaling. **E**. PDGF signaling. **F**. WNT and non-canonical WNT signaling. **G**. FGF signaling with MyoFB-Mat cells as the dominant FGF9 signaling source and MyoFB-Int and MyoFB-Mat cells as the dominant FGF18 signaling source in mesenchymal populations. FGF10 signaling from MatFB to epithelial cells.

### Functional properties of mesenchymal cell types in the neonatal lung

Functional properties of mesenchymal cell populations marked by *Fgf18^CreERT2^; ROSA^TDTomato^* and *Pdgfra^EGFP^*were examined in *in vitro* cultures of sorted C1, C2, and C3 cells from mice induced with tamoxifen from P2-P5 and harvested on P7 (Figs. 2A, 7A). Consistent with the cell sorting parameters, following one day *in vitro* culture (DIV1), C1 and C2 cells expressed GFP and were negative for TDTomato expression, while C3 cells expressed both GFP and TDTomato (Fig. 7B). Violin plots of GFP expression from scRNA seq data showed GFP in all cell samples with highest levels in samples C2 (Fig. 7C). Consistent with sorting and DIV1 culture, TDTomato was highly expressed in C3 cells with only trace amounts detected in samples C1 and C2 (Fig. 7D).

**Figure 7.**
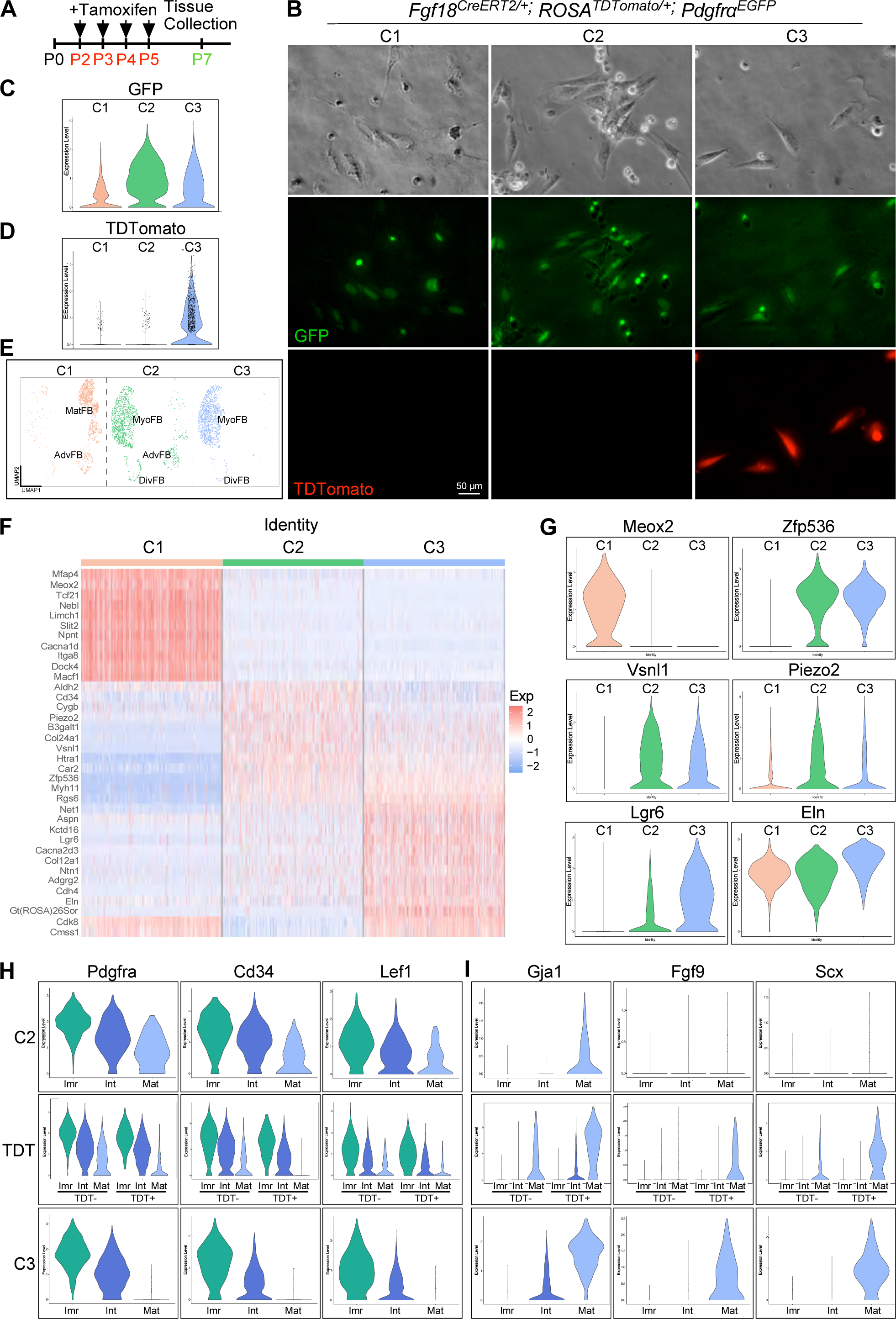
Gene expression patterns of sorted populations of lung mesenchymal cells. **A**. Experimental scheme for lineage labeling of mesenchymal cell populations in *Fgf18^CreER^; ROSA^TDTomato^*; *Pdgfra^EGFP^* mice. Black arrowheads indicate tamoxifen injection from P2-P5. P7 lung tissue was then dissociated, sorted for GFP and TDTomato, and cultured. **B**. Cultures of C1 (TDTomato^−^, GFP^low^), C2 (TDTomato^−^, GFP^High^) and C3 (TDTomato^+^, GFP^High^) cells after one day *in vitro* culture (DIV1) were imaged with brightfield optics and fluorescence for *Pdgfra^EGFP^* (GFP) and FGF18 (TDT). **C**. Violin plots derived from scRNA seq data, grouped by sample, showing highest GFP expression in C2 cells and lower expression in C1 and C3 cells. **D**. Violin plot, grouped by sample, showing highest TDTomato expression in C3 and trace expression in C1 and C2 cells. **E**. UMAP showing the distribution of C1, C2 and C3 cells. C1 cells include 2.8% of MyoFB, 10.8% of DivFB, 53.4% of AdvFB, and 94.3% of MatFB; C2 cells include 43.8% of MyoFB, 61.9% of DivFB, 45.5% of AdvFB, and 5.1% of MatFB; C3 cells include 53.3% of MyoFB, 27.3% of DivFB, 1.1% of AdvFB, and 0.6% of MatFB. **F**. Heatmap showing genes enriched in C1, C2, and C3 cells. **G**. Violin plots showing expression of genes enriched in C1, C2, and C3 cells. **H, I**. Violin plots comparing the MyoFB expression of immature markers *Pdgfra, Cd34*, and *Lef1* (**H**), and mature markers, *Gja1, Fgf9*, and *Scx* (**i**) in C2 cells, TDTomato negative cells (TDT^−^), TDTomato positive cells (TDT^+^), and C3 cells.

Gene signatures that define the sorted C1, C2 and C3 cells are consistent with C1 containing MatFBs, a small subcluster of AdvFB (vascular smooth muscle) and a very small number of DivFB (Fig. 7E, Supplemental Fig. 2). C1 cells were highly enriched for markers such as *Meox2, Tcf21*, and *Nebl* (Fig. 7F, G). Sample C2 cells were represented by all MyoFB clusters and were the predominant contributor to DivFB and AdvFB clusters (Fig. 7E, Supplemental Fig. 2). C2 cells were enriched for MyoFB-Imr markers such as *Vsnl* and *Piezo2* (Figs. 2D, 7F, G). C3 cells contain primarily MyoFB and a small proportion of DivFB (Fig. 7E, Supplemental Fig. 2). C3 cells were enriched for markers expressed in MyoFB-Mat cells such as *Lgr6, Cdh4*, and *Eln* (Fig. 7F, G). C2 and C3 cells were more related to each other and are distinguished from C1 by the expression of common markers such as the transcription factor *Zfp536*, and MyoFB genes such at *Tagln*, *Myh11* and *Itga8*. Although C2 and C3 cells show significant similarities, they are distinguished by expression of *FGF18 (TDT)* and other genes such as *Piezo2*, high in C2, and *Col12a1*, high in C3 (Fig. 7D, F).

Further comparison of MyoFB in C2 and C3 cells show that genes associated with less mature cells such as *Pdgfra, Cd34,* and *Lef1* were expressed at higher levels in C2 cells with a profile that was most similar to that of TDTomato negative cells (Fig. 7H, Supplemental Table 1E). By contrast, C3 cells expressed these genes at relatively lower levels with a profile that was most similar to TDTomato positive cells (Fig. 7H, Supplemental Table 1E). Similarly, genes associated with more mature cells such as *Gja1, Fgf9*, and *Scx*, were expressed at relatively lower levels in C2 cells with a profile that was most similar to that of TDTomato negative cells. By contrast, C3 cells expressed these genes at relatively higher levels with a profile that was most similar to TDTomato positive cells (Fig. 7I). Overlaying TDTomato expression onto the MyoFB UMAP shows relatively low expression in MyoFB-Imr cells and higher expression in MyoFB-Int and MyoFB-Mat cells (Supplemental Fig. 8).

Trajectory, diffusion, and velocity analysis predicted that MyoFB-Imr cells transition to MyoFB-Int cells and finally to MyoFB-Mat cells (Fig. 5A-E). To explore this prediction, C2 cells were maintained in culture and monitored for GFP and TDTomato expression (Fig. 8A, B). Although negative for TDTomato expression on DIV1, TDTomato expressing cells became visible by DIV2 and increased by DIV4. Sorting DIV4 cells identified TDTomato^−^, GFP^Low^ (C1*); TDTomato^−^, GFP^High^ (C2*); and TDTomato^+^, GFP^High^ (C3*) cells (Fig. 8C). Of the GFP expressing cells at DIV4, 47% were GFP^Low^, 38.6% were GFP^High^, and 14.3% activated TDTomato expression (C3* cells) (Fig. 8D). TDTomato^−^, GFP^−^ cells presumably are derived from expansion of contaminating GFP^−^ cells in the original sorted sample or transition of GFP^+^ C2 cells to a GFP^−^ cell.

**Figure 8.**
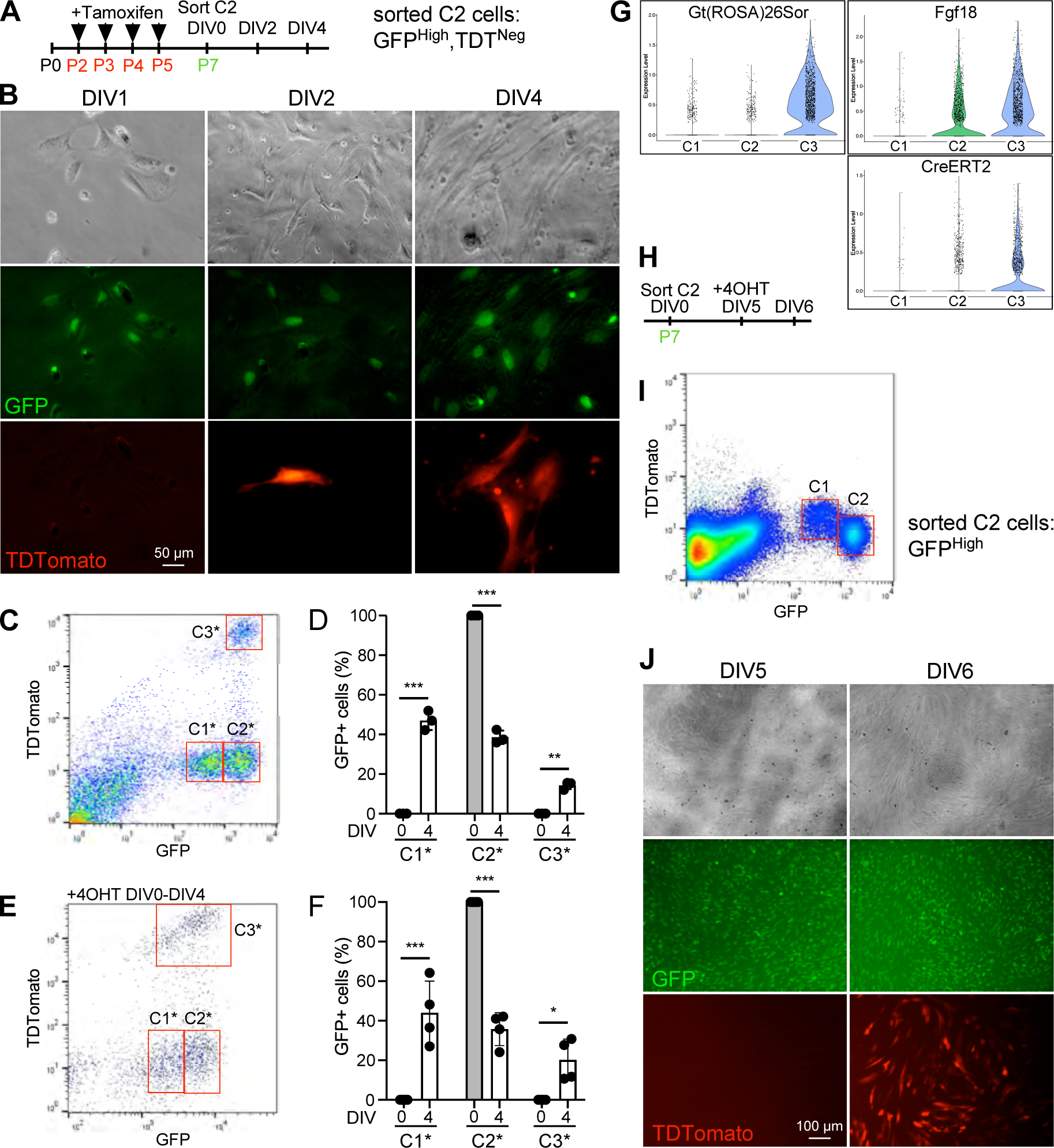
Spontaneous maturation of C2 cells *in vitro* is marked by expression of the endogenous and targeted ROSA26 locus. **A**. Experimental schematic. Black arrow heads indicate tamoxifen injection from P2 to P5. At P7, C2 (TDTomato^−^, GFP^High^) cells were plated on Matrigel-coated plates. **B**. Brightfield and fluorescent images of C2 cells at DIV1, DIV2 and DIV4. At DIV1 only TDTomato^−^, GFP^+^ cells are present. At DIV2, some cells become TDTomato^+^ and the number of TDTomato^+^ cells increase by DIV4. **C-F**. Quantification of C2 cell maturation in the absence (**C, D**) or presence (**E, F**) of 4-OHT. **C, E**. At DIV4, C2 cell cultures were sorted for GFP and TDTomato expressing cells, identifying three populations of cells labeled as C1* (GFP^Low^), C2* (GFP^High^), and C3*(TDT^+^, GFP^High^). **D, F**. Quantification of the percentage of C1*, C2* and C3* cells at DIV0 (grey bars) and DIV4 (open bars) (**D**, n=3; **F**, n=4). 4-OHT does not significantly change percentages of C1*, C2* and C3* cells at DIV4. **G**. Violin plots derived from scRNA seq data showing expression of the endogenous ROSA26 long non-coding RNA (Gt(ROSA)26Sor), *Fgf18*, and transgenic *Fgf18^CreERT2^* (CreERT2) in C1, C2, and C3 cells. **h, i**. Experimental schema for sorting and culturing GFP^Low^ and GFP^High^ (C2 + C3) cells from P7 *Fgf18^CreERT2^; ROSA^TDTomato^; Pdgfra^EGFP^* lungs from mice that were not induced with tamoxifen. **j**. C2 cells (GFP^High^) cultured in the absence of 4-OHT for five days. Imaging shows abundant GFP^pos^ cells and no TDTomato expressing cells on DIV5. Adding 4-OHT to the culture medium on DIV5 results in abundant TDTomato positive cells on DIV6. Statistics: 2-way ANOVA with Fisher’s LSD test; n=3 or 4; ns, not significant; * *P* < 0.05, ** *P* < 0.001, *** *P* < 0.0001.

Interestingly, the activation of TDTomato expression (derived from the *ROSA^TDTomato^* reporter) in culture occurred in the absence of tamoxifen, suggesting that expression of the ROSA26 gene locus was itself regulated and is a reporter for maturation of immature MyoFB. To control for this, we cultured C2 cells in the presence of 4-Hydroxytamoxifen (4-OHT). However, even in the presence of 4-OHT, similar numbers of TDTomato^−^, GFP^Low^ (C1*); TDTomato^−^, GFP^High^ (C2*); and TDTomato^+^, GFP^High^ (C3*) cells were present at DIV4 (Fig. 8E, F), demonstrating that additional recombination at the ROSA26 locus is not occurring during culture.

Consistent with regulation of the ROSA26 locus, expression of the endogenous ROSA26 long non-coding RNA (Gt(ROSA)26Sor) in the scRNA seq data was present at low levels in C1 and C2 cells and high levels in C3 cells (Fig. 8G), similar to the expression of TDTomato in C1, C2 and C3 cells (Fig. 7D). Furthermore, expression of endogenous *Fgf18* and transgenic *Fgf18^CreERT2^*were detected in C2 cells (Fig. 8G). The expression of CreERT2 in C2 cells suggests that in mice induced with tamoxifen, *ROSA^TDTomato^* recombination had occurred but the *ROSA26* allele and TDTomato were expressed at very low or undetectable levels. To control for any leaky CreERT2 activity, GFP^High^ cells, which should contain the equivalent of both C2 and C3 cells, were sorted from *Fgf18^CreERT2^; ROSA^TDTomato^*; *Pdgfra^EGFP^* mice that were not induced with tamoxifen (Fig. 8H, I). Sorted cells, harvested at P7, were then cultured for five days (Fig. 8J). At DIV5, cultured GFP^High^ cells continued to express GFP but did not express TDTomato. However, when these DIV5 cultures were treated with 4-OHT for 24 hr, TDTomato activity was robustly detected at DIV6, demonstrating that these cells expressed *Fgf18^CreERT2^,* that CreER was not leaky and could be induced *in vitro*, and that the *ROSA26* locus was active in these cells after 5 days in culture.

C1 cells (TDTomato^−^, GFP^low^) are primarily composed of MatFB and AdvFB (Fig. 7E). Trajectory, diffusion, and velocity analysis shows potential differentiation pathways within these two populations (Fig. 5A, F-I); however, there were no trajectories linking MatFB/AdvFB and MyoFB. This is consistent with 3H-thymidine uptake studies which demonstrated that lipid containing interstitial cells (corresponding to MatFB) were distinct from non-lipid containing interstitial cells (corresponding to MyoFB) (Brody and Kaplan, 1983). To determine if C1 cells have the capacity to generate TDTomato^+^ MyoFB, similar to what we observed with C2 cells (TDTomato^−^, GFP^High^), sorted C1 cells were maintained in culture for four days. At DIV4, C1 cells were larger and more spread out, the intensity of GFP fluorescence was decreased, but no TDTomato^+^ cells were observed (Fig. 9A). Cell sorting confirmed the stability of the C1 population showing that of GFP-expressing cells, over 98% of cells were in the GFP^Low^ sorting gate (C1*), 1.3% of cells were GFP^High^ (C2*), and no cells had detectable TDTomato (C3*) (Fig. 9B, C). Culture of C1 cells in the presence of 4-OHT, followed by sorting, showed a small percentage of TDTomato^−^, GFP^High^ C2* cells (11±7%) and TDTomato^+^, GFP^High^ C3* cells (5±5%) cells at DIV4. It is likely that these cells arise from proliferation of contaminating MyoFB progenitors (GFP^High^) within the sorted C1 cells as the sorting gates are adjacent to each other, but we cannot rule out differentiation of a GFP^Low^ MatFB or AdvFB into a GFP^High^ MyoFB progenitor.

**Figure 9.**
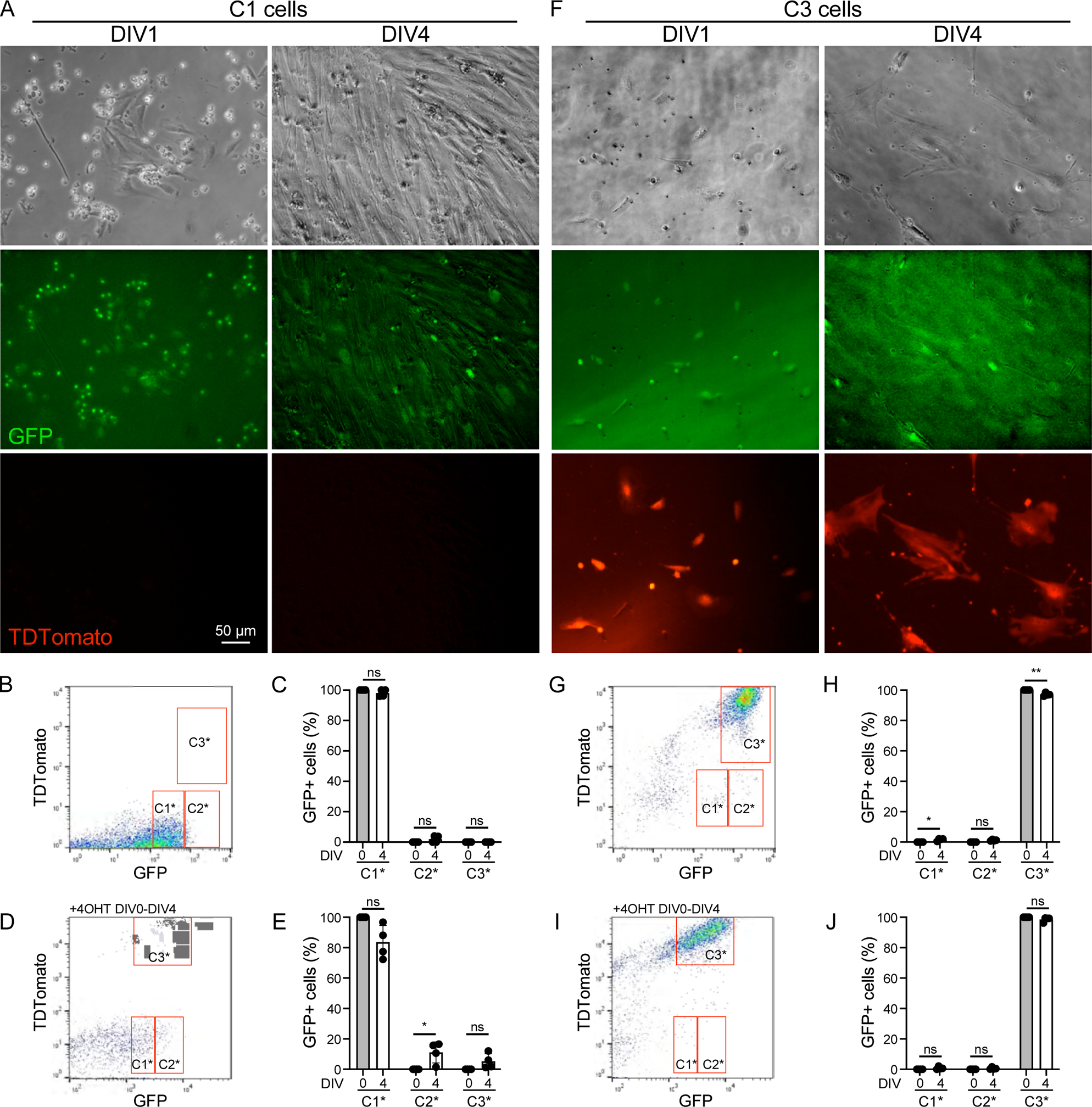
MatFB and TDTomato expressing MyoFB remain stable in *in vitro* culture. **A**. Brightfield and fluorescent images of C1 cells at DIV1 and DIV4. At both DIV1 and DIV4 only GFP^+^, TDTomato^−^ cells are present. **B-E**. Quantification of C1 cell maturation in the absence (**B, C**) or presence (**D, E**) of 4-OHT. **B, D**. At DIV4, C1 cell cultures were sorted for GFP and TDTomato expressing cells. Of the GFP expressing cells, most retained the GFP^Low^ (C1*) phenotype. **C, E**. Quantification of the percentage of C1*, C2* and C3* cells at DIV0 (grey bars) and DIV4 (open bars) (**C**, n=3; **E**, n=4). 4-OHT does not significantly change percentages of C1*, C2* and C3* cells at DIV4. **F**. Brightfield and fluorescent images of C3 cell cultures at DIV1 and DIV4. At both DIV1 and DIV4 TDTomato^+^, GFP^+^ cells are present. **G-J**. Quantification of C3 cell maturation in the absence (**G, H**) or presence (**I, J**) of 4-OHT. **G, I**. At DIV4, C3 cell cultures were sorted for GFP and TDTomato expressing cells. Of the GFP expressing cells, most retained the TDTomato^+^, GFP^High^ (C3*) phenotype. **H, J**. Quantification of the percentage of C1*, C2* and C3* cells at DIV0 (grey bars) and DIV4 (open bars) (**H**, n=3; **J**, n=4). 4-OHT does not significantly change percentages of C1*, C2* and C3* cells at DIV4. Statistics: 2-way ANOVA with Fisher’s LSD test; n=4; ns, not significant; * *P* < 0.05, ** *P* < 0.001.

C3 cells (TDTomato^+^, GFP^High^) are composed almost exclusively of MyoFB and proliferating cells with a MyoFB gene expression signature (Fig. 7E, Supplemental Fig. 2). Following four days in culture, C3 cells increased in size and retained TDTomato expression (Fig. 9F). Cell sorting confirmed the stability of the C3 population showing that of GFP-expressing cells, over 96% also expressed TDTomato (C3*), 2.1% were in the GFP^Low^ sorting gate (C1*) and 1.3% of cells were in the GFP^High^; TDTomato^−^ sorting gate (C2*) (Fig. 9G, H). Culture of C3 cells in the presence of 4-OHT did not affect the numbers of TDTomato^−^, GFP^Low^ (C1*); TDTomato^−^, GFP^High^ (C2*); or TDTomato^+^, GFP^High^ (C3*) cells present at DIV4 (Fig. 9I, J).

## Discussion

Alveologenesis is a unique stage of lung development where the myofibroblast (MyoFB), an essential cell that guides the formation of secondary septa, is a transient cell population that is only present during the primary phase of alveologenesis (Branchfield et al., 2016; Duong et al., 2022; Gao et al., 2022; Hagan et al., 2020; Li et al., 2015; McGowan et al., 2008; Narvaez Del Pilar et al., 2022; Riccetti et al., 2020; Yamada et al., 2005; Zepp et al., 2021). The MyoFB, also called a secondary crest myofibroblast (SCMF), is first observed in the mouse around P2-P3, at the initiation of alveologenesis, and most can no longer be detected by the end of primary alveologenesis, ∼P14-P15. By contrast, adventitial fibroblasts (AdvFB) and matrix fibroblasts (MatFB), which include lipofibroblasts, are present during alveologenesis and persist in the adult lung. Although embryonic mesenchymal cells that give rise to lung mesenchymal lineages have been identified (Li et al., 2015; Li et al., 2018; Moiseenko et al., 2017; Yamada et al., 2005), the progenitor cell that allows expansion of the MyoFB population during primary alveologenesis has yet to be identified.

The MyoFB produces elastin, and its intrinsic contractile properties are essential for secondary septation (Li et al., 2019; Li et al., 2015; Lindahl et al., 1997; Zepp et al., 2021). The MyoFB has previously been identified and isolated based on its expression of high levels of GFP driven by the *Pdgfra* gene promoter (Endale et al., 2017; Kimani et al., 2009; McGowan and McCoy, 2014; Zepp et al., 2021). The *Pdgfra-*GFP^High^ MyoFB was also found to express *Fgf18*, a member of the fibroblast growth factor family, and *Fgf18* expression in the lung is greatly increased during the primary phase of alveologenesis in rodents and humans (Boucherat et al., 2007; Chailley-Heu et al., 2005; Franco-Montoya et al., 2011; McGowan and McCoy, 2015; Ruiz-Camp and Morty, 2015). This expression pattern was confirmed by lineage labeling with an *Fgf18^CreER^* allele crossed to the *ROSA^TDTomato^*lineage reporter – FGF18 (TDT) (Hagan et al., 2019; Hagan et al., 2020).

In an effort to further characterize lung mesenchymal cell populations during alveologenesis, cells from dissociated P7 lung tissue were sorted based on expression of GFP (*Pdgfra^EGFP^*) and TDTomato (*Fgf18^CreER^; ROSA^TDTomato^*) following four days of tamoxifen administration from P2-P5. This configuration allowed separation of the *Pdgfra-*GFP^High^ cell population into two groups, TDTomato^−^, GFP^High^ (C2 cells) and TDTomato^+^, GFP^High^ (C3 cells). Single cell sequencing revealed that both of these cell populations contained MyoFB and that the TDTomato^−^ cells also contained AdvFB. Interestingly, the GFP^High^, TDTomato^−^ cell population was enriched for proliferating cells with a MyoFB gene signature and the GFP^High^, TDTomato^+^ cells were enriched for markers of mature MyoFB. In culture, some of the GFP^High^, TDTomato^−^ cells were able to produce GFP^High^, TDTomato^+^ cells, suggesting that the initial C2 cell population contains a proliferating progenitor that gives rise to mature myofibroblasts. Activation of TDTomato expression during this maturation process occurred in the absence of tamoxifen and thus indicates that upregulation of *ROSA^TDTomato^*gene expression is not induced but is a regulated event that marks MyoFB maturation. Consistent with this, single cell sequencing data shows both increased TDTomato and increased endogenous *ROSA26* long non-coding RNA expression in C3 MyoFB compared to C2 MyoFB.

In human postnatal day 1 lung, Duong et al. identified two types of MyoFB (Duong et al., 2022). MyoFB_1 was thought to be a progenitor based on higher expression of *PDGFRA*, *IGF1*, and *MEF2A,* and MyoFB_2 was thought to represent a more mature population with elevated expression of *ACTA2, MYH11,* and *MYOCD*. In our data, in MyoFB-Imr cells, we see higher levels of *Pdgfra*, slightly elevated *Mef2a* but lower levels of *Igf1,* and in MyoFB-Mat cells we see elevated *Myocd*, but no difference in *Acta2* or *Myh11* levels. These differences may reflect differences between mouse and human or differences in the relative developmental stage.

A unique feature of the MyoFB-Mat population is the expression of *Gja1* (Cx43)*, Fgf9*, and *Scx.* Immunostaining showed that these proteins or their lineage reporters are present in alveolar myofibroblasts in septal tip locations and alveolar ducts. The function of these genes in the neonatal lung is not known; however, in other tissues, both *Gja1* and *Scx* have been shown to maintain a myofibroblast phenotype or promote a fibroblast to myofibroblast transition (Bagchi et al., 2016; Paw et al., 2017). The MyoFB-Mat cells also express periostin (*Postn*), a direct transcriptional target of SCX (Nagalingam et al., 2022). The function of FGF9 in the neonatal lung is also not known, but in the adult, it may function to inhibit human IPF fibroblast differentiation to myofibroblasts and promote a migratory phenotype (Joannes et al., 2016). We posit that during alveologenesis, FGF9, along with FGF18, could function to regulate migration or maturation of MyoFB-Imr progenitor cells, which, relative to MyoFB-Mat cells, express higher levels of *Fgfr1* and *Fgfr2*. Unlike FGF18, FGF9 can also signal to lung epithelial cells via FGFR3 (Yin et al., 2013; Yin and Ornitz, 2020; Yin et al., 2016), and thus could coordinate MyoFB-Mat-epithelial interactions.

MyoFB-Imr cells are likely to contain a progenitor population that gives rise to MyoFB-Int and MyoFB-Mat cells. Consistent with this model, MyoFB-Imr cells are enriched for *CD34*, *Eng* (Edoglin, CD105), *Lef1* (lymphoid enhancer factor 1), *Stc1* (Stanniocalcin-1), and *Plcb1* (phospholipase C, beta 1), genes associated with a progenitor phenotype in multiple tissues and cell lineages (Hennrick et al., 2007; Schelker et al., 2021; Sidney et al., 2014; Song and Kim, 2022). In adult lung and kidney fibrosis, lineage tracing showed that pericytes could give rise to pathological myofibroblasts (Armulik et al., 2011; Barron et al., 2016; Humphreys et al., 2010; Hung et al., 2013); however, in the lung this remains controversial and one lineage tracing study showed that pericytes are excluded as a major source of pathological myofibroblasts (Rock et al., 2011). Interestingly, *Pdgfrb*, an accepted marker for pericytes, is enriched in MyoFB-Imr cells compared to MyoFB-Int and MyoFB-Mat cells; however, MyoFB-Imr cells express low levels of other pericyte markers, *Cspg4* (*Ng2*) and *Foxj1*, suggesting that they are not closely related to pericytes.

The mechanisms that regulate the disappearance of MyoFB after primary alveologenesis are still elusive. Fang et al. showed that in the absence of epithelial Wnt secretion, ⍺SMA^+^ cells persisted within alveolar septa (Fang et al., 2022); however, they attributed this to increased endothelial to mesenchymal transition rather than failure to lose preexisting MyoFB. MyoFB-Imr cells, and at lower levels, MyoFB-Mat cells, express the canonical WNT receptors LRP6 and FZD1. By contrast, expression of the non-canonical WNT ligand, *Wnt5a*, is increased in MyofB-Int and MyoFB-Mat cells. WNT5a functions to suppress canonical WNT/β-catenin signaling and induce caspase activity in differentiating embryonic stem cells (Bisson et al., 2015). *Vsnl1* (Visinin-like 1), encoding a WNT/β-catenin-regulated calcium sensing protein, is expressed at high levels in MyoFB-Imr and MyoFB-Int cells and low levels in MyoFB-Mat cells. In colorectal cancer, VSNL1 functions to suppress apoptosis (Tage et al., 2023). By analogy, its down regulation in MyoFB-Mat cells could contribute to their eventual loss. Future studies will be needed to determine if WNT signaling directly regulates MyoFB maturation and loss of WNT signaling triggers MyoFB apoptosis at the end of primary alveologenesis.

Although we focus on the MyoFB lineage in this study, our scRNA seq data revealed unique subclusters of MatFB and AdvFB and cell culture shows that MatFB (the predominant cell type in C1 cells) are stable and do not spontaneously convert to TDTomato^+^ cells.

Bronchovascular bundles are conduits for conducting airways and associated vasculature and contain adventitial or bronchovasular cuff fibroblasts (Dahlgren et al., 2019; Narvaez Del Pilar et al., 2022; Sun et al., 2022; Travaglini et al., 2020; Tsukui et al., 2020). Subclustering of AdvFB and MatFB-Im,Cuff cells identified two subclusters that are related to bronchovascular cuff fibroblasts. We have termed these cells as Bronchovascular Cuff FB Type 1 and Type 2 (BVC-Type 1, BVC-Type2). Both of these clusters express common marker such as *Dner, Igtbl1, Mecom, Serpinf1,* and *Twist2*. These clusters are distinguished by expression of *Adh7, Lgr5*, and *Wif1* in BVC-Type 1 cells and *Col14a1,* and *Pi16* in BVC-Type2 cells. This expression pattern suggests that BVC-Type 1 cells may have progenitor-like properties as *Lgr5* expression is a marker of stem cells and the Wnt inhibitor, WIF1, can regulate progenitor cell proliferation (Barker et al., 2007; Yu et al., 2022). Interestingly, Bisson et al. have identified an adult *Lgr5^+^* lung mesenchymal cell population that localizes to the alveolar region and promotes alveolar differentiation via activation of WNT signaling (Lee et al., 2017). BVC-Type 2 cells, which express *Col14a1,* may have a greater role in producing ECM components. *Col14a1^+^* cells have been localized to perivascular regions of developing lung (Mizikova et al., 2022).

Meox2 is expressed in MatFB and AdvFB and is excluded from smooth muscle. Consistent with the expression patterns shown by Narvaez del Pilar et al.(Narvaez Del Pilar et al., 2022), our immunostaining shows MEOX2^+^ cells localized to bronchovascular cuff regions adjacent to peribronchial and perivascular smooth muscle cells and co-localizing with cells that express low levels of *Pdgfra^EGFP^*. The expression pattern of *Meox2* is similar to that of *Tcf21* suggesting that these genes could be co-regulated.

### Limitations

These studies have only analyzed mesenchymal cells that express *Pdgfra^EGFP^* and lineage traced *Fgf18^CreERT2^*cells at a single time point (P7) near the peak of alveologenesis. However, since alveologenesis occurs dynamically over time and likely progresses from proximal to distal as the lung grows, any time point should contain all the required cellular components. All cells analyzed in this study are heterozygous for *Fgf18* and *Pdgfra*. Although no overt phenotypes have been associated with haploinsufficiency of these genes, given their importance for mesenchymal cell differentiation, we cannot rule out subtle effects on relative cluster size and quantitative levels of gene expression. In addition to *Pdgfra^EGFP^*cells, there are likely other mesenchymal cells present in the neonatal lung that do not express *Pdgfra^EGFP^*. The identity of these *Pdgfra^EGFP^* negative cells will require high resolution single cell analysis of the whole lung or other lineage markers that can be used for cell sorting and sequencing. In this study we only sequenced one sample for each sorted population. However, each sample was composed of pooled cells from three mice, to minimize variability among animals. Finally, in Table 1 we have tried to cross reference the names given to the multitude of mesenchymal cell types that have been identified by many investigators both in the neonate and adult. We apologize if we have inadvertently missed some names given to mesenchymal cell types.

## Materials and Methods

### Animals

All mice (*Mus musculus*) were housed in a pathogen-free barrier facility. All studies were performed under a protocol approved by the Institutional Animal Care at Washington University in Saint Louis (Approval No. 20190110). Mice of both sexes were used. Mice were maintained on a mixed FVB, C57BL/6J, 129X1/SvJ genetic background. Mouse strains: *Fgf18^CreERT2^* (*Fgf18^tm2.2(cre/ERT2)Dor^*) (Hagan et al., 2019), *ROSA^TDTomato^* (*Gt*(*ROSA*)*26Sor^tm9(CAG-TDTomato)Hze/J)^* (Madisen et al., 2010), *Pdgfra^EGFP^* (*Pdgfra^tm11(EGFP)Sor/J^*) (Hamilton et al., 2003), *Fgf9^βGal^* (Huh et al., 2015), *Scx^Cre^*(Sugimoto et al., 2013). The day of birth was assigned as P0.

### Tamoxifen administration

Tamoxifen was administered to neonates by intraperitoneal injection at a dose of 150 µg/mouse/day on four sequential days beginning at P2. Tamoxifen (Sigma; T5648) was prepared at a concentration of 15 µg/µl dissolved in corn oil or sunflower seed oil.

### Tissue preparation for CLARITY and immunohistochemistry

At designated collection times, mice were sacrificed with an overdose of a cocktail containing ketamine and xylazine, and perfused with PBS through the right ventricle. For immunohistochemistry the lungs were fixed via intratracheal inflation with 4% paraformaldehyde (PFA) (Electron Microscopy Sciences; 15714-S) at a pressure of 20 cm H_2_O. Lungs were post-fixed overnight at 4°C with gentle agitation. For immunohistochemistry, samples were washed three times in PBS for 20 min, cut into lobes, dehydrated in ethanol and xylene, embedded in paraffin, sectioned (5 μm), and stained with hematoxylin and eosin (H&E) or used for immunostaining.

For CLARITY the lungs were fixed in 0.5 ml of 4% PFA by intratracheal injection with a 26g needle. Lungs were post-fixed overnight at 4°C in 4% PFA with gentle agitation and then washed 5 times with PBS for 20 min at room temperature.

### Antibodies and chemical reagents

The following chemical reagent and antibodies were used for immunofluorescence: Alexa Fluor 633 hydrazide (Elastin) (A30634, Invitrogen), goat anti-TDTomato (1:200, MBS448092, MyBioSource.com), rabbit anti-Connexin 43 (Cx43) (1:200, SC-9059, Santa Cruz Biotechnology), mouse anti-Smooth Muscle Actin (αSMA) (1:300, Clone 1A4, Dako), goat anti-TAGLN (1:200, ab10135, Abcam), chicken anti-β Galactosidase (1:200, ab9361, Abcam), FITC-conjugated mouse anti-CD34 (1:200, 11-0341-82, Invitrogen), mouse anti-LEF1 (1:100, SC-374522, Santa Cruz Biotechnology), rabbit anti-PDGFRA (1:100, sc-338, Santa Cruz Biotechnology), chicken anti-GFP (1:200, AB-2307313, Aveslab), rabbit anti-MEOX2 (1:200, NBP2-30647, Novus Biological). Alexa-488, 555, 594 or Alexa-647-conjugated donkey anti-goat, anti-rabbit, anti-mouse, anti-chicken secondary antibodies (all from Invitrogen; A11055, A21432, A21447, A21206, A21207, A31573, A21202, A31570, A21203, A31571, A78952), Alexa-488 or Alexa 594-conjugated donkey anti-chicken secondary antibodies (703-545-155, and 703-585-155 from Jackson ImmunoResearch).

### Tissue clearing (CLARITY)

Samples were optically cleared using the SHIELD tissue preservation approach (Park et al., 2018) and subsequently delipidated using ETC active CLARITY. Briefly, 4% PFA-fixed tissues were immersed in SHIELD-off polyepoxy solution for 3 days at 4°C and subsequently polymerized by immersion in SHIELD-on solution at 37°C for 24 hr. Preserved samples were delipidated in a 4% sodium dodecyl sulfate buffer for 24 hr at 45°C, and were transferred to a SmartBatch+ electrophoretic tissue clearing (ETC) platform (LifeCanvas Technologies, Boston, MA), and cleared for 30 hr at 35V, 1250mA at 42°C. Following clearing, samples were washed in PBS for 24 hr. Lungs were washed 3 times with PBS/0.2% Triton X-100 for 1 hr at 37°C and stained for elastin with Alexa Fluor 633 Hydrazide (Shen et al., 2012) (1:1000, Thermo Fisher Scientific; A30634) in PBS/0.2% Triton X-100 for 24 hr at 37°C. After staining, the lungs were washed 4 times with PBS/0.2% Triton X-100 for 2 hr at 37°C, and then stored in PBS/0.2% Triton X-100 at 4°C. Prior to imaging, the refractive index of lungs were matched to 1.52 using Easy Index (LifeCanvas Technologies).

### Confocal imaging and image analysis

High resolution imaging datasets of cleared lungs were acquired on a Zeiss LSM 980 microscope using the Airyscan 2 SR-4Y mode with either a 40x/1.2 water immersion or a 63x/1.4 oil immersion objective. Raw Airyscan files were processed using Huygens Professional (Scientific Volume Imaging, Hilversum, Netherlands) and visualized in Imaris (Oxford Instruments). Thin section confocal images were acquired on a Zeiss LSM 980 microscope with a 40x/1.2 water immersion or a 63x/1.4 oil immersion objective and visualized in Imaris.

### Flow cytometry

Lung cells were isolated as described (Hagan et al., 2020) with minor modifications. *Fgf18^CreERT2/+^*; *ROSA^TDTomato/+^*; *Pdgfra^EGFP/+^*pups were injected with tamoxifen (150 µg) on P2, P3, P4, P5 and anesthetized and sacrificed on P7 with an overdose of a cocktail containing ketamine and xylazine. The lungs were dissected and placed in PBS on ice, minced with a razor blade with 50 µl of dispase (5,000 Caseinolytic Units, Discovery Labware, Inc. 354235), and then transferred to PBS containing 833 units/ml dispase, 0.01 mg/ml DNase I (Roche; 10104159001) and 2 mg/ml collagenase/dispase (Roche, 10269638) at 37°C for 40 min with gentle agitation. Digestion enzymes were then inactivated with FACS buffer (10% FBS, 1 mM EDTA in PBS), samples were sequentially filtered through a 70 µM cell strainer (Thermo Fisher Scientific; 22363548), a 40 µM cell strainer (Thermo Fisher Scientific; 22363547), and then centrifuged at 800 x g for 4 min. Red blood cells were lysed with ACK lysis buffer (Gibco; A10492-01) and cells were resuspended in FACS buffer. SYTOX blue (1 µl, Thermo Fisher Scientific, S34857) was added to each tube for live/dead sorting. Cells were sorted using a SY3200 “Synergy” cell sorter (Sony Biotechnology) or a Beckman Coulter MoFlo cell sorter (Beckman Coulter Life Sciences) with a 100 µm nozzle directly into FACS buffer or culture media (1.5 ml) on ice. Four cell populations were collected: TDTomato^+^, GFP^−^; TDTomato^+^, GFP^High^; TDTomato^−^, GFP^High^; and TDTomato^−^, GFP^Low^. Single channel controls (*Fgf18^CreERT2^*; *ROSA^TDTomato^*; *Pdgfra^EGFP^* mice) were used to set compensation at the time of sorting and a fluorescence minus one control (*ROSA^TDTomato^* lacking *Fgf18^CreERT2^*) was used as a negative control. Plotted percentage of the four populations of cells was calculated relative to the total number of sorted cells. Freshly sorted cells were either placed into culture or submitted to the Washington University Genome Technology Access Center for scRNA seq.

### Cell culture

Sorted cells were resuspended in DMEM/F12 (Gibco, 11330-032) with 20% Fetal Bovine Serum (Gibco, 26140-079), 2% penicillin/streptomycin, and plated on Matrigel (1:2 dilution)-coated 24 or 48 wells plates and incubated at 37°C in 5% CO_2_. Media was changed daily during the course of the experiment. Cells were imaged using a Leica DM IL LED inverted microscope (Leica Microsystems, CMS GmbH).

### Single cell RNA sequencing (scRNA seq)

For single cell sequencing, C1, C2, and C3 cells (Fig. 2A) were sorted from three individual mice. Each of the three samples (C1, C2, and C3) were pooled from the three mice and were used for droplet-based scRNA seq using 10x Genomics 3’v3 kits at the Washington University Genome Technology Access Center (GTAC). Flow sorted populations were loaded on individual lanes in a Chromium Controller as per manufacturer instructions. Following their preparation per the 10x Genomics protocol, each library was sequenced using an Illumina NovaSeq 6000 instrument. After demultiplexing of Illumina runs, FASTQ files were processed by the 10x Genomics Cell Ranger pipeline (version **7.0.0**) with exonic and intronic reads incorporated in the final molecule counts.

#### Cell clustering and annotation

For initial clustering, cells from *Fgf18* heterozygous (*Fgf18^CreERT2/+^*; *ROSA^TDTomato/+^*; *Pdgfra^EGFP/+^*) P7 mice were clustered with cells obtained from *Fgf18* conditional knockout (*Fgf18^CreERT2/f^*; *ROSA^TDTomato/+^*; *Pdgfra^EGFP/+^*) P7 mice obtained with an identical flow sorting approach. Expression matrices were input into Seurat for clustering analysis. Low quality cells were filtered by feature counts (>2300 genes/cell, <6500 genes/cell) and mitochondrial gene content (< 10%). Following merge, batch effect correction was performed with the *Harmony* package (Korsunsky et al., 2019). After Louvain clustering and UMAP visualization (using Harmony embeddings), cell type-specific markers were identified with Seurat’s *FindAllMarkers* function, with log fold change set to 0.25. Annotation for broad cell types (myofibroblasts, matrix and adventitial fibroblasts, dividing cells) was performed using known markers, and specific marker genes were identified for subtypes within these categories. Rare cells identified as undesired cell types (epithelium, endothelium, immune) based on known marker genes were removed after clustering. For all downstream analyses, a subset of *Fgf18* heterozygous cells from the combined object was used.

#### Trajectory analysis of mesenchymal cells

For trajectory analysis, spliced and unspliced mRNAs for all genes in all cells were determined using Velocyto (https://github.com/velocyto-team/velocyto.py) using a cell sorted BAM file for each sample as input. The resulting loom files were input to scVelo, merged in anndata object, and velocities were estimated using scv.tl.velocity. For visualization, dimensional reduction was performed using the standard scanpy (https://scanpy.readthedocs.io; v1.9.1) pipeline with PCA and UMAP, and cluster assignments annotated based on the Seurat analysis above. To calculate diffusion maps, we built the k-nearest neighbor graph (k = 25) using the first 30 principal components in scanpy and used the scanpy.tl.diffmap function to build diffusion maps with an exponential kernel (method=”umap”). For projection of velocities onto diffusion maps, DM coordinates were added to adata.obs in the appropriate subsetted adata objects. The R Bioconductor package *slingshot* (https://bioconductor.org/packages/release/bioc/html/slingshot.html) was used to calculate the lineage structure and pseudotime, which were projected onto the same diffusion maps.

#### Cell Chat analysis

For receptor-ligand analysis, a publicly available P7 mouse lung scRNA seq filtered expression matrix was used (sample GSM4504963) (Zepp et al., 2021). After initial clustering and annotation with Seurat, mesenchymal cell populations were removed from these data and then *merge* was used to combined our annotated fibroblast populations (with dividing populations removed). The combined object was then normalized, re-clustered using the Seurat pipeline as above, and inputted into the CellChat package in R (Jin et al., 2021). Default settings were used, and for *filterCommunication*, min.cells was set to 10.

### Reverse Transcriptase Quantitative PCR

RNA was isolated from sorted TDTomato^+^ cells according to manufacturer’s instructions using the Arcturus PicoPure RNA Isolation Kit (Applied biosystems, Thermo Fisher Scientific, 12204-01). P7 whole lung and adult heart RNA, isolated with Trizol Reagent (Ambion by life technologies, 15596026) served as positive and negative control, respectively, for gene expression. cDNA was generated by reverse transcription of up to 9 ng RNA from each sample using iScript Reverse Transcription Supermix for RT-qPCR (BioRad, 1708841). mRNA expression was determined on a StepOnePlus^TM^ Real-Time PCR System (Thermo Fisher, 4376600) using TaqMan^TM^ Fast Advanced Master Mix (Thermo Fisher, 4444557) and TaqMan assay probes (*Gapdh*, Thermo Fisher, Mm99999915_g1; *Pdpn*, Thermo Fisher, Mm01348912_g1; *Wt1*, Thermo Fisher, Mm00460570_m1). Relative gene expression was calculated based on the average cycle (Ct) value of technical triplicates, normalized to *Gapdh* control, and reported as fold change (2^−ΔΔCT^).

### Statistical analysis

Data were analyzed with Prism software using analysis of variance (ANOVA). For quantification of gene expression by qPCR, 3-4 individual samples were measured and analyzed using one-way ANOVA with Tukey’s multiple comparisons test. For quantification of sorted cell populations 3-4 independent cultures derived from different animals were analyzed using 2-way ANOVA with Fishers Least Significant Difference (LSD) test for multiple comparisons. The data are reported as the means ± SD, and changes with P values less than 0.05 were considered to be statistically significant.

## Supporting information

Genomic data tables

## Acknowledgments

We thank H. McNeill for providing Scx-Cre mice and L. Li for technical assistance.

## Author contributions

Y.Y., J.R.K., A.S.H., and D.M.O. conceptualized and designed the study; Y.Y., J.R.K., A.S.H., D.P., S.D., P.B., and D.M.O. developed methodology; Y.Y., J.R.K., D.P., S.D., P.B., D.M.O. performed experiments; Y.Y., J.R.K., D.P., S.D., P.B., and D.M.O. analyzed data; Y.Y., J.R.K., and D.M.O. drafted manuscript; Y.Y., J.R.K., D.P., S.D., P.B., A.S.H., and D.M.O. reviewed and edited manuscript; Y.Y., J.R.K., and D.M.O. acquired funding.

## Supplemental Figure Legends

**Supplemental Movie 1. 3D rendering of a confocal image of whole mount cleared lung from a P7 mouse.** 3D image of a lung from tamoxifen induced *Fgf18^CreER^; ROSA^TDTomato^* (Red), *Pdgfra^EGFP^* (Green) mouse, stained for elastin with Alexa Fluor 633 Hydrazide (white).

**Supplemental Figure 1. Quantitative RT-PCR of RNA from sorted T cells (TDTomato^+^, GFP^−^), whole lung, and whole heart.** T cells express the mesothelial marker Wilms tumor 1 homolog (*Wt1*) and the AT1 cell marker Podoplanin (*Pdpn, T1⍺*). *Wt1* was detected at low levels in whole lung and not detected in heart. *Pdpn* was expressed in whole lung but not in heart. n=3-4, ANOVA with Tukey’s multiple comparison, ** *P* < 0.002, *** *P* < 0.0001.

**Supplemental Figure 2. Analysis of cell cycle stages and distribution in sorted mesenchymal groups. A**. S phase cells, determined by Seurat Cell-Cycle Scoring are enriched in C2 cells and are present at low levels in C3 cells. **B**. G2/M phase cells are enriched in C2 cells and are present at low levels in C3 cells. **C**. reference UMAP showing primary cell types. **D**. Relative numbers of proliferating cells within primary sorted cell types normalized to the number of cells in each sample.

**Supplemental Figure 3. Identification of immature and mature myofibroblast subtypes. A**. UMAP and violin plots comparing myofibroblast subclusters in our data set with markers identified by Narvaez Del Pilar et al. that were present in both neonatal and adult lung (*Cdh4, Hhip*, and *Lgr6*) and *Pdgfra*-high expressing cells that were only present in neonatal lung. In our data set, *Cdh4, Hhip*, and *Lgr6* were associated with MyoFB-Mat cells and *Pdgfra^EGFP^* (GFP) and endogenous *Pdgfra-high* and *Pdgfrb* were associated with MyoFB-Imr cells. **B**. Genes associated with progenitor cells, *Cd34* and *Lef1*, were enriched in MyoFB-Imr cells whereas *Aspn, Fgf9,* and *Gja1* were enriched in MyoFB-Mat cells. *Fgf18* lineage TDTomato was enriched in MyoFB-Int and MyoFB-Mat cells.

**Supplemental Figure 4. Expression patterns of genes in the proximal and distal P7 mouse lung. A**. In the distal lung, FGF9 (βGal)^+^, LEF1^−^ (red arrow), FGF9 (βGal)^+^, LEF1^+^ (yellow arow), and FGF9 (βGal)^−^, LEF1^+^ (green arrow) cells were identified, corresponding to MyoFB-Mat, MyoFB-Int, and MyoFB-Imr cells, respectively. **B**. Cx43^−^, LEF1^+^ cells (green arrow) and Cx43^+^, LEF1^+^ cells (yellow arrow) correspond to MyoFB-Imr and MyoFB-Int cells, respectively. **C**. SCX (TDT)^+^, LEF1^−^, MKI67^−^ cells (red arrows) and SCX (TDT)^+^, LEF1^+^, MKI67^+^ cells (green arrows), correspond to MyoFB-Mat and MyoFB-Imr cells, respectively. **D**. FGF18 (TDT), *Pdgfra^EGFP^* (GFP)^High^ cells correspond with MyoFB-Mat cells (red arrows); FGF18 (TDT)^Low^, *Pdgfra^EGFP^*(GFP)^Low^ cells correspond with MyoFB-Imr cells (green arrow); FGF18 (TDT)^−^, *Pdgfra^EGFP^* (GFP), Cx43^+^ cells correspond with MatFB (white arrow). **E**. Validation of endogenous PDGFRA and *Pdgfra^EGFP^* (GFP) expression. Dashed box in the left image indicates area shown at higher magnification in the following images. *Pdgfra^EGFP^*(GFP)^+^ nuclear staining is localized in cells that have cytoplasmic staining for PDGFRA. *Pdgfra^EGFP^* (GFP)^Low^, ⍺SMA^+^ cells are marked as MyoFB-Mat (red arrows); *Pdgfra^EGFP^* (GFP)^High^, ⍺SMA^+^ cells correspond to MyoFB-Imr cells (green arrows).

**Supplemental Figure 5. Gene Ontologies for differentially expressed genes in mesenchymal cell types. A-D**. GO analysis of DEGs for MyoFB-Imr (**A**), MyoFB (TDTomato^−^) (**B**), MyoFB-Mat (**C**), and MyoFB (TDTomato^+^) (**D**). MyoFB (excluding proliferating cells) grouped by subcluster or grouped by TDTomato expression were used to identify DEGs using the FindAllMarkers function in Seurat (Log2FV > 0.25). **E**. MatFB (including subclusters Synthetic, AF1-Type 1, and AF1-Type 2) were used to identify DEGs using the FindAllMarkers function in Seurat (Log2FV > 0.25). **F**. AdvFB (including AdvFB subclusters 0, 1, 2 and BVC-Type 1) were used to identify DEGs using the FindAllMarkers function in Seurat (Log2FV > 0.25).

**Supplemental Figure 6. Subclustering of MatFB-Im,Cuff cells. A.** UMAP showing markers unique to MatFB-Im,Cuff cells and markers expressed in both MatFB-Im,Cuff and AdvFB cells. **B.** UMAP shows representative genes that are selectively expressed in subclusters of MatFB-Im,Cuff cells. Subcluster 0, designated MatFB-0 (synthetic), are enrichment for ribosomal and ECM genes and express high levels of *Nebl* and *Nav2*. Subcluster 1, designated BVC-Type 1, express *Adh7*, *Pdgfrb, Twist2, Dner* and *Col14a1* and corresponds to a subtype of an adult adventitial or bronchovascular cuff fibroblast. Subcluster 2 (Synthetic/Stress) expressed stress response factors such as *Fos*.

**Supplemental Figure 7. Subclustering of AdvFB cells. A.** UMAP showing markers unique to AdvFB cells. **B**. UMAP shows representative genes that are selectively expressed in subclusters. Subclusters 0 is enriched for expression of *Hhip, Boc,* and *Lsamp* and subcluster 1 is enriched for expression of *Zfp36* and *Icam1.* Subcluster 2, designated BVC-Type 2, express *Pi16* and *Ebf2* and corresponds to a subtype of adult lung adventitial fibroblasts. Subcluster 3 (VSM, pericyte) express *Acta2, Myh11, Tagln, Thsd4, Sost, Cbr2, Rgs5,* and *Cspg4* and is likely composed of vascular smooth muscle cells, a cell similar to vascular mural cells, and pericytes. Subcluster 4 expressed *Grm7*, *Pcks5, Musk,* and corresponds to an undefined lung stromal cell.

**Supplemental Figure 8. Expression level of TDTomato correlates with maturation of MyoFB**. UMAP of MyoFB clusters is plotted in the XY axis and FGF18 (TDT) expression is plotted in the Z axis.

## References

Adams, T. S., Schupp, J. C., Poli, S., Ayaub, E. A., Neumark, N., Ahangari, F., Chu, S. G., Raby, B. A., DeIuliis, G., Januszyk, M., et al. (2020). Single-cell RNA-seq reveals ectopic and aberrant lung-resident cell populations in idiopathic pulmonary fibrosis. Sci Adv 6, eaba1983.

Ahn, S. and Joyner, A. L. (2004). Dynamic changes in the response of cells to positive hedgehog signaling during mouse limb patterning. Cell 118, 505–516.

Amy, R. W., Bowes, D., Burri, P. H., Haines, J. and Thurlbeck, W. M. (1977). Postnatal growth of the mouse lung. J Anat 124, 131–151.

Armulik, A., Genove, G. and Betsholtz, C. (2011). Pericytes: developmental, physiological, and pathological perspectives, problems, and promises. Dev Cell 21, 193–215.

Bagchi, R. A., Roche, P., Aroutiounova, N., Espira, L., Abrenica, B., Schweitzer, R. and Czubryt, M. P. (2016). The transcription factor scleraxis is a critical regulator of cardiac fibroblast phenotype. BMC Biol 14, 21.

Bancalari, E. and Jain, D. (2019). Bronchopulmonary Dysplasia: 50 Years after the Original Description. Neonatology 115, 384–391.

Barker, N., van Es, J. H., Kuipers, J., Kujala, P., van den Born, M., Cozijnsen, M., Haegebarth, A., Korving, J., Begthel, H., Peters, P. J., et al. (2007). Identification of stem cells in small intestine and colon by marker gene Lgr5. Nature 449, 1003–1007.

Barron, L., Gharib, S. A. and Duffield, J. S. (2016). Lung Pericytes and Resident Fibroblasts: Busy Multitaskers. Am J Pathol 186, 2519–2531.

Bisson, J. A., Mills, B., Paul Helt, J. C., Zwaka, T. P. and Cohen, E. D. (2015). Wnt5a and Wnt11 inhibit the canonical Wnt pathway and promote cardiac progenitor development via the Caspase-dependent degradation of AKT. Dev Biol 398, 80–96.

Bose, B. and Shenoy, P. S. (2016). Aging induced loss of stemness with concomitant gain of myogenic properties of a pure population of CD34(+)/CD45(−) muscle derived stem cells. Int J Biochem Cell Biol 70, 1–12.

Bostrom, H., Gritli-Linde, A. and Betsholtz, C. (2002). PDGF-A/PDGF alpha-receptor signaling is required for lung growth and the formation of alveoli but not for early lung branching morphogenesis. Dev Dyn 223, 155–162.

Boucherat, O., Benachi, A., Barlier-Mur, A. M., Franco-Montoya, M. L., Martinovic, J., Thebaud, B., Chailley-Heu, B. and Bourbon, J. R. (2007). Decreased lung fibroblast growth factor 18 and elastin in human congenital diaphragmatic hernia and animal models. Am J Respir Crit Care Med 175, 1066–1077.

Branchfield, K., Li, R., Lungova, V., Verheyden, J. M., McCulley, D. and Sun, X. (2016). A three-dimensional study of alveologenesis in mouse lung. Dev Biol 409, 429–441.

Brody, J. S. and Kaplan, N. B. (1983). Proliferation of alveolar interstitial cells during postnatal lung growth. Evidence for two distinct populations of pulmonary fibroblasts. Am Rev Respir Dis 127, 763–770.

Chailley-Heu, B., Boucherat, O., Barlier-Mur, A. M. and Bourbon, J. R. (2005). FGF-18 is upregulated in the postnatal rat lung and enhances elastogenesis in myofibroblasts. Am J Physiol Lung Cell Mol Physiol 288, L43–51.

Chao, C. M., Moiseenko, A., Zimmer, K. P. and Bellusci, S. (2016). Alveologenesis: key cellular players and fibroblast growth factor 10 signaling. Mol Cell Pediatr 3, 17.

Dahlgren, M. W., Jones, S. W., Cautivo, K. M., Dubinin, A., Ortiz-Carpena, J. F., Farhat, S., Yu, K. S., Lee, K., Wang, C., Molofsky, A. V., et al. (2019). Adventitial Stromal Cells Define Group 2 Innate Lymphoid Cell Tissue Niches. Immunity 50, 707–722 e706.

Diaz-Flores, L., Gutierrez, R., Garcia, M. P., Saez, F. J., Diaz-Flores, L., Jr., Valladares, F. and Madrid, J. F. (2014). CD34+ stromal cells/fibroblasts/fibrocytes/telocytes as a tissue reserve and a principal source of mesenchymal cells. Location, morphology, function and role in pathology. Histol Histopathol 29, 831–870.

Dorry, S. J., Ansbro, B. O., Ornitz, D. M., Mutlu, G. M. and Guzy, R. D. (2020). FGFR2 Is Required for AEC2 Homeostasis and Survival after Bleomycin-induced Lung Injury. Am J Respir Cell Mol Biol 62, 608–621.

Du, Y., Guo, M., Whitsett, J. A. and Xu, Y. (2015). ’LungGENS’: a web-based tool for mapping single-cell gene expression in the developing lung. Thorax 70, 1092–1094.

Du, Y., Kitzmiller, J. A., Sridharan, A., Perl, A. K., Bridges, J. P., Misra, R. S., Pryhuber, G. S., Mariani, T. J., Bhattacharya, S., Guo, M., et al. (2017). Lung Gene Expression Analysis (LGEA): an integrative web portal for comprehensive gene expression data analysis in lung development. Thorax 72, 481–484.

Duong, T. E., Wu, Y., Sos, B. C., Dong, W., Limaye, S., Rivier, L. H., Myers, G., Hagood, J. S. and Zhang, K. (2022). A single-cell regulatory map of postnatal lung alveologenesis in humans and mice. Cell Genom 2.

El Agha, E., Herold, S., Al Alam, D., Quantius, J., MacKenzie, B., Carraro, G., Moiseenko, A., Chao, C. M., Minoo, P., Seeger, W., et al. (2014). Fgf10-positive cells represent a progenitor cell population during lung development and postnatally. Development 141, 296–306.

Endale, M., Ahlfeld, S., Bao, E., Chen, X., Green, J., Bess, Z., Weirauch, M. T., Xu, Y. and Perl, A. K. (2017). Temporal, spatial, and phenotypical changes of PDGFRalpha expressing fibroblasts during late lung development. Dev Biol 425, 161–175.

Fang, Y., Shao, H., Wu, Q., Wong, N. C., Tsong, N., Sime, P. J. and Que, J. (2022). Epithelial Wntless regulates postnatal alveologenesis. Development 149.

Franco-Montoya, M. L., Boucherat, O., Thibault, C., Chailley-Heu, B., Incitti, R., Delacourt, C. and Bourbon, J. R. (2011). Profiling target genes of FGF18 in the postnatal mouse lung: possible relevance for alveolar development. Physiol Genomics 43, 1226–1240.

Gao, F., Li, C., Danopoulos, S., Al Alam, D., Peinado, N., Webster, S., Borok, Z., Kohbodi, G. A., Bellusci, S. and Minoo, P. (2022). Hedgehog-responsive PDGFRa(+) fibroblasts maintain a unique pool of alveolar epithelial progenitor cells during alveologenesis. Cell Rep 39, 110608.

Green, J., Endale, M., Auer, H. and Perl, A. K. (2016). Diversity of Interstitial Lung Fibroblasts Is Regulated by Platelet-Derived Growth Factor Receptor alpha Kinase Activity. Am J Respir Cell Mol Biol 54, 532–545.

Guo, M., Morley, M. P., Jiang, C., Wu, Y., Li, G., Du, Y., Zhao, S., Wagner, A., Cakar, A. C., Kouril, M., et al. (2023). Guided construction of single cell reference for human and mouse lung. Nat Commun 14, 4566.

Hagan, A. S., Boylan, M., Smith, C., Perez-Santamarina, E., Kowalska, K., Hung, I. H., Lewis, R. M., Hajihosseini, M. K., Lewandoski, M. and Ornitz, D. M. (2019). Generation and validation of novel conditional flox and inducible Cre alleles targeting fibroblast growth factor 18 (Fgf18). Dev Dyn 248, 882–893.

Hagan, A. S., Zhang, B. and Ornitz, D. M. (2020). Identification of a FGF18-expressing alveolar myofibroblast that is developmentally cleared during alveologenesis. Development 147, dev.181032.

Haghverdi, L., Buettner, F. and Theis, F. J. (2015). Diffusion maps for high-dimensional single-cell analysis of differentiation data. Bioinformatics 31, 2989–2998.

Hamilton, T. G., Klinghoffer, R. A., Corrin, P. D. and Soriano, P. (2003). Evolutionary divergence of platelet-derived growth factor alpha receptor signaling mechanisms. Mol Cell Biol 23, 4013–4025.

He, J., Duan, H., Xiong, Y., Zhang, W., Zhou, G., Cao, Y. and Liu, W. (2013). Participation of CD34-enriched mouse adipose cells in hair morphogenesis. Mol Med Rep 7, 1111–1116.

Hennrick, K. T., Keeton, A. G., Nanua, S., Kijek, T. G., Goldsmith, A. M., Sajjan, U. S., Bentley, J. K., Lama, V. N., Moore, B. B., Schumacher, R. E., et al. (2007). Lung cells from neonates show a mesenchymal stem cell phenotype. Am J Respir Crit Care Med 175, 1158–1164.

Huh, S. H., Warchol, M. E. and Ornitz, D. M. (2015). Cochlear progenitor number is controlled through mesenchymal FGF receptor signaling. Elife 4, 1–17.

Humphreys, B. D., Lin, S. L., Kobayashi, A., Hudson, T. E., Nowlin, B. T., Bonventre, J. V., Valerius, M. T., McMahon, A. P. and Duffield, J. S. (2010). Fate tracing reveals the pericyte and not epithelial origin of myofibroblasts in kidney fibrosis. Am J Pathol 176, 85–97.

Hung, C., Linn, G., Chow, Y. H., Kobayashi, A., Mittelsteadt, K., Altemeier, W. A., Gharib, S. A., Schnapp, L. M. and Duffield, J. S. (2013). Role of lung pericytes and resident fibroblasts in the pathogenesis of pulmonary fibrosis. Am J Respir Crit Care Med 188, 820–830.

Jin, S., Guerrero-Juarez, C. F., Zhang, L., Chang, I., Ramos, R., Kuan, C. H., Myung, P., Plikus, M. V. and Nie, Q. (2021). Inference and analysis of cell-cell communication using CellChat. Nat Commun 12, 1088.

Joannes, A., Brayer, S., Besnard, V., Marchal-Somme, J., Jaillet, M., Mordant, P., Mal, H., Borie, R., Crestani, B. and Mailleux, A. A. (2016). FGF9 and FGF18 in idiopathic pulmonary fibrosis promote survival and migration and inhibit myofibroblast differentiation of human lung fibroblasts in vitro. Am J Physiol Lung Cell Mol Physiol 310, L615–629.

Kimani, P. W., Holmes, A. J., Grossmann, R. E. and McGowan, S. E. (2009). PDGF-Ralpha gene expression predicts proliferation, but PDGF-A suppresses transdifferentiation of neonatal mouse lung myofibroblasts. Respir Res 10, 119.

Koenitzer, J. R., Wu, H., Atkinson, J. J., Brody, S. L. and Humphreys, B. D. (2020). Single-Nucleus RNA-Sequencing Profiling of Mouse Lung. Reduced Dissociation Bias and Improved Rare Cell-Type Detection Compared with Single-Cell RNA Sequencing. Am J Respir Cell Mol Biol 63, 739–747.

Korsunsky, I., Millard, N., Fan, J., Slowikowski, K., Zhang, F., Wei, K., Baglaenko, Y., Brenner, M., Loh, P. R. and Raychaudhuri, S. (2019). Fast, sensitive and accurate integration of single-cell data with Harmony. Nat Methods 16, 1289–1296.

La Manno, G., Soldatov, R., Zeisel, A., Braun, E., Hochgerner, H., Petukhov, V., Lidschreiber, K., Kastriti, M. E., Lonnerberg, P., Furlan, A., et al. (2018). RNA velocity of single cells. Nature 560, 494–498.

Lee, J. H., Tammela, T., Hofree, M., Choi, J., Marjanovic, N. D., Han, S., Canner, D., Wu, K., Paschini, M., Bhang, D. H., et al. (2017). Anatomically and Functionally Distinct Lung Mesenchymal Populations Marked by Lgr5 and Lgr6. Cell 170, 1149–1163 e1112.

Li, C., Lee, M. K., Gao, F., Webster, S., Di, H., Duan, J., Yang, C. Y., Bhopal, N., Peinado, N., Pryhuber, G., et al. (2019). Secondary crest myofibroblast PDGFRalpha controls the elastogenesis pathway via a secondary tier of signaling networks during alveologenesis. Development 146.

Li, C., Li, M., Li, S., Xing, Y., Yang, C. Y., Li, A., Borok, Z., De Langhe, S. and Minoo, P. (2015). Progenitors of secondary crest myofibroblasts are developmentally committed in early lung mesoderm. Stem Cells 33, 999–1012.

Li, R., Bernau, K., Sandbo, N., Gu, J., Preissl, S. and Sun, X. (2018). Pdgfra marks a cellular lineage with distinct contributions to myofibroblasts in lung maturation and injury response. Elife 7.

Liberti, D. C., Kremp, M. M., Liberti, W. A., 3rd, Penkala, I. J., Li, S., Zhou, S. and Morrisey, E. E. (2021). Alveolar epithelial cell fate is maintained in a spatially restricted manner to promote lung regeneration after acute injury. Cell Rep 35, 109092.

Ligresti, G., Raslan, A. A., Hong, J., Caporarello, N., Confalonieri, M. and Huang, S. K. (2023). Mesenchymal cells in the Lung: Evolving concepts and their role in fibrosis. Gene 859, 147142.

Lindahl, P., Karlsson, L., Hellstrom, M., Gebre-Medhin, S., Willetts, K., Heath, J. K. and Betsholtz, C. (1997). Alveogenesis failure in PDGF-A-deficient mice is coupled to lack of distal spreading of alveolar smooth muscle cell progenitors during lung development. Development 124, 3943–3953.

Madisen, L., Zwingman, T. A., Sunkin, S. M., Oh, S. W., Zariwala, H. A., Gu, H., Ng, L. L., Palmiter, R. D., Hawrylycz, M. J., Jones, A. R., et al. (2010). A robust and high-throughput Cre reporting and characterization system for the whole mouse brain. Nat Neurosci 13, 133–140.

McGowan, S. E., Grossmann, R. E., Kimani, P. W. and Holmes, A. J. (2008). Platelet-derived growth factor receptor-alpha-expressing cells localize to the alveolar entry ring and have characteristics of myofibroblasts during pulmonary alveolar septal formation. Anat Rec (Hoboken) 291, 1649–1661.

McGowan, S. E. and McCoy, D. M. (2014). Regulation of fibroblast lipid storage and myofibroblast phenotypes during alveolar septation in mice. Am J Physiol Lung Cell Mol Physiol 307, L618–631.

McGowan, S. E. and McCoy, D. M. (2015). Fibroblast growth factor signaling in myofibroblasts differs from lipofibroblasts during alveolar septation in mice. Am J Physiol Lung Cell Mol Physiol 309, L463–474.

McGowan, S. E. and Torday, J. S. (1997). The pulmonary lipofibroblast (lipid interstitial cell) and its contributions to alveolar development. Annu Rev Physiol 59, 43–62.

Mizikova, I., Lesage, F., Cyr-Depauw, C., Cook, D. P., Hurskainen, M., Hanninen, S. M., Vadivel, A., Bardin, P., Zhong, S., Carpen, O., et al. (2022). Single-Cell RNA Sequencing-Based Characterization of Resident Lung Mesenchymal Stromal Cells in Bronchopulmonary Dysplasia. Stem Cells 40, 479–492.

Moiseenko, A., Kheirollahi, V., Chao, C. M., Ahmadvand, N., Quantius, J., Wilhelm, J., Herold, S., Ahlbrecht, K., Morty, R. E., Rizvanov, A. A., et al. (2017). Origin and characterization of alpha smooth muscle actin-positive cells during murine lung development. Stem Cells 35, 1566–1578.

Mougin, Z., Huguet Herrero, J., Boileau, C. and Le Goff, C. (2021). ADAMTS Proteins and Vascular Remodeling in Aortic Aneurysms. Biomolecules 12.

Muhl, L., Genove, G., Leptidis, S., Liu, J., He, L., Mocci, G., Sun, Y., Gustafsson, S., Buyandelger, B., Chivukula, I. V., et al. (2020). Single-cell analysis uncovers fibroblast heterogeneity and criteria for fibroblast and mural cell identification and discrimination. Nat Commun 11, 3953.

Mund, S. I., Stampanoni, M. and Schittny, J. C. (2008). Developmental alveolarization of the mouse lung. Dev Dyn 237, 2108–2116.

Muzumdar, M. D., Tasic, B., Miyamichi, K., Li, L. and Luo, L. (2007). A global double-fluorescent Cre reporter mouse. Genesis 45, 593–605.

Nabhan, A. N., Brownfield, D. G., Harbury, P. B., Krasnow, M. A. and Desai, T. J. (2018). Single-cell Wnt signaling niches maintain stemness of alveolar type 2 cells. Science 359, 1118–1123.

Nagalingam, R. S., Chattopadhyaya, S., Al-Hattab, D. S., Cheung, D. Y. C., Schwartz, L. Y., Jana, S., Aroutiounova, N., Ledingham, D. A., Moffatt, T. L., Landry, N. M., et al. (2022). Scleraxis and fibrosis in the pressure-overloaded heart. Eur Heart J 43, 4739–4750.

Narvaez Del Pilar, O., Gacha Garay, M. J. and Chen, J. (2022). Three-axis classification of mouse lung mesenchymal cells reveals two populations of myofibroblasts. Development 149.

Park, J., Ivey, M. J., Deana, Y., Riggsbee, K. L., Sorensen, E., Schwabl, V., Sjoberg, C., Hjertberg, T., Park, G. Y., Swonger, J. M., et al. (2019). The Tcf21 lineage constitutes the lung lipofibroblast population. Am J Physiol Lung Cell Mol Physiol 316, L872–L885.

Park, Y. G., Sohn, C. H., Chen, R., McCue, M., Yun, D. H., Drummond, G. T., Ku, T., Evans, N. B., Oak, H. C., Trieu, W., et al. (2018). Protection of tissue physicochemical properties using polyfunctional crosslinkers. Nat Biotechnol.

Paw, M., Borek, I., Wnuk, D., Ryszawy, D., Piwowarczyk, K., Kmiotek, K., Wojcik-Pszczola, K. A., Pierzchalska, M., Madeja, Z., Sanak, M., et al. (2017). Connexin43 Controls the Myofibroblastic Differentiation of Bronchial Fibroblasts from Patients with Asthma. Am J Respir Cell Mol Biol 57, 100–110.

Riccetti, M., Gokey, J. J., Aronow, B. and Perl, A. T. (2020). The elephant in the lung: Integrating lineage-tracing, molecular markers, and single cell sequencing data to identify distinct fibroblast populations during lung development and regeneration. Matrix Biol 91-92, 51–74.

Rippa, A. L., Alpeeva, E. V., Vasiliev, A. V. and Vorotelyak, E. A. (2021). Alveologenesis: What Governs Secondary Septa Formation. Int J Mol Sci 22.

Rock, J. R., Barkauskas, C. E., Cronce, M. J., Xue, Y., Harris, J. R., Liang, J., Noble, P. W. and Hogan, B. L. (2011). Multiple stromal populations contribute to pulmonary fibrosis without evidence for epithelial to mesenchymal transition. Proc Natl Acad Sci U S A 108, E1475–1483.

Ruiz-Camp, J. and Morty, R. E. (2015). Divergent fibroblast growth factor signaling pathways in lung fibroblast subsets: where do we go from here? Am J Physiol Lung Cell Mol Physiol 309, L751–755.

Sanger, C. S., Cernakova, M., Wietecha, M. S., Garau Paganella, L., Labouesse, C., Dudaryeva, O. Y., Roubaty, C., Stumpe, M., Mazza, E., Tibbitt, M. W., et al. (2023). Serine protease 35 regulates the fibroblast matrisome in response to hyperosmotic stress. Sci Adv 9, eadh9219.

Schelker, R. C., Kratzer, A., Muller, G., Brochhausen, C., Hart, C., Stempfl, T., Heudobler, D., Moehle, C., Herr, W., Iberl, S., et al. (2021). Stanniocalcin 1 is overexpressed in multipotent mesenchymal stromal cells from acute myeloid leukemia patients. Hematology 26, 565–576.

Schmidt, K. L., Marcus-Gueret, N., Adeleye, A., Webber, J., Baillie, D. and Stringham, E. G. (2009). The cell migration molecule UNC-53/NAV2 is linked to the ARP2/3 complex by ABI-1. Development 136, 563–574.

Shah, P. S., Sankaran, K., Aziz, K., Allen, A. C., Seshia, M., Ohlsson, A., Lee, S. K. and Canadian Neonatal, N. (2012). Outcomes of preterm infants <29 weeks gestation over 10-year period in Canada: a cause for concern? J Perinatol 32, 132–138.

Shen, Z., Lu, Z., Chhatbar, P. Y., O’Herron, P. and Kara, P. (2012). An artery-specific fluorescent dye for studying neurovascular coupling. Nat Methods 9, 273–276.

Sidney, L. E., Branch, M. J., Dunphy, S. E., Dua, H. S. and Hopkinson, A. (2014). Concise review: evidence for CD34 as a common marker for diverse progenitors. Stem Cells 32, 1380–1389.

Song, B. and Kim, C. H. (2022). Cell-autonomous PLCbeta1 modulation of neural stem/progenitor cell proliferation during adult hippocampal neurogenesis. Neurosci Lett 791, 136899.

Stoll, B. J., Hansen, N. I., Bell, E. F., Walsh, M. C., Carlo, W. A., Shankaran, S., Laptook, A. R., Sanchez, P. J., Van Meurs, K. P., Wyckoff, M., et al. (2015). Trends in Care Practices, Morbidity, and Mortality of Extremely Preterm Neonates, 1993-2012. JAMA 314, 1039–1051.

Street, K., Risso, D., Fletcher, R. B., Das, D., Ngai, J., Yosef, N., Purdom, E. and Dudoit, S. (2018). Slingshot: cell lineage and pseudotime inference for single-cell transcriptomics. BMC Genomics 19, 477.

Sugimoto, Y., Takimoto, A., Hiraki, Y. and Shukunami, C. (2013). Generation and characterization of ScxCre transgenic mice. Genesis 51, 275–283.

Sun, K. H., Chang, Y., Reed, N. I. and Sheppard, D. (2016). alpha-Smooth muscle actin is an inconsistent marker of fibroblasts responsible for force-dependent TGFbeta activation or collagen production across multiple models of organ fibrosis. Am J Physiol Lung Cell Mol Physiol 310, L824–836.

Sun, X., Perl, A. K., Li, R., Bell, S. M., Sajti, E., Kalinichenko, V. V., Kalin, T. V., Misra, R. S., Deshmukh, H., Clair, G., et al. (2022). A census of the lung: CellCards from LungMAP. Dev Cell 57, 112–145 e112.

Tabula Muris, C., Overall, c., Logistical, c., Organ, c., processing, Library, p., sequencing, Computational data, a., Cell type, a., Writing, g., et al. (2018). Single-cell transcriptomics of 20 mouse organs creates a Tabula Muris. Nature 562, 367–372.

Tage, H., Yamaguchi, K., Nakagawa, S., Kasuga, S., Takane, K., Furukawa, Y. and Ikenoue, T. (2023). Visinin-like 1, a novel target gene of the Wnt/beta-catenin signaling pathway, is involved in apoptosis resistance in colorectal cancer. Cancer Med 12, 13426–13437.

Tirosh, I., Izar, B., Prakadan, S. M., Wadsworth, M. H., 2nd, Treacy, D., Trombetta, J. J., Rotem, A., Rodman, C., Lian, C., Murphy, G., et al. (2016). Dissecting the multicellular ecosystem of metastatic melanoma by single-cell RNA-seq. Science 352, 189–196.

Travaglini, K. J., Nabhan, A. N., Penland, L., Sinha, R., Gillich, A., Sit, R. V., Chang, S., Conley, S. D., Mori, Y., Seita, J., et al. (2020). A molecular cell atlas of the human lung from single-cell RNA sequencing. Nature 587, 619–625.

Tsukui, T., Sun, K. H., Wetter, J. B., Wilson-Kanamori, J. R., Hazelwood, L. A., Henderson, N. C., Adams, T. S., Schupp, J. C., Poli, S. D., Rosas, I. O., et al. (2020). Collagen-producing lung cell atlas identifies multiple subsets with distinct localization and relevance to fibrosis. Nat Commun 11, 1920.

Vila Ellis, L. and Chen, J. (2021). A cell-centric view of lung alveologenesis. Dev Dyn 250, 482–496.

Whitsett, J. A. and Weaver, T. E. (2015). Alveolar development and disease. Am J Respir Cell Mol Biol 53, 1–7.

Xie, T., Wang, Y., Deng, N., Huang, G., Taghavifar, F., Geng, Y., Liu, N., Kulur, V., Yao, C., Chen, P., et al. (2018). Single-Cell Deconvolution of Fibroblast Heterogeneity in Mouse Pulmonary Fibrosis. Cell Rep 22, 3625–3640.

Yamada, M., Kurihara, H., Kinoshita, K. and Sakai, T. (2005). Temporal expression of alpha-smooth muscle actin and drebrin in septal interstitial cells during alveolar maturation. J Histochem Cytochem 53, 735–744.

Yang, J., Hernandez, B. J., Martinez Alanis, D., Narvaez del Pilar,O., Vila-Ellis, L., Akiyama, H., Evans, S. E., Ostrin, E. J. and Chen, J. (2016). The development and plasticity of alveolar type 1 cells. Development 143, 54–65.

Yin, Y., Betsuyaku, T., Garbow, J. R., Miao, J., Govindan, R. and Ornitz, D. M. (2013). Rapid induction of lung adenocarcinoma by fibroblast growth factor 9 signaling through FGF receptor 3. Cancer Res 73, 5730–5741.

Yin, Y. and Ornitz, D. M. (2020). FGF9 and FGF10 activate distinct signaling pathways to direct lung epithelial specification and branching. Sci Signal 13, eaay4353.

Yin, Y., Ren, X., Smith, C., Guo, Q., Malabunga, M., Guernah, I., Zhang, Y., Shen, J., Sun, H., Chehab, N., et al. (2016). Inhibition of fibroblast growth factor receptor 3-dependent lung adenocarcinoma with a human monoclonal antibody. Dis Model Mech 9, 563–571.

Yu, Q. C., Geng, A., Preusch, C. B., Chen, Y., Peng, G., Xu, Y., Jia, Y., Miao, Y., Xue, H., Gao, D., et al. (2022). Activation of Wnt/beta-catenin signaling by Zeb1 in endothelial progenitors induces vascular quiescence entry. Cell Rep 41, 111694.

Yuan, T., Volckaert, T., Redente, E. F., Hopkins, S., Klinkhammer, K., Wasnick, R., Chao, C. M., Yuan, J., Zhang, J. S., Yao, C., et al. (2019). FGF10-FGFR2B Signaling Generates Basal Cells and Drives Alveolar Epithelial Regeneration by Bronchial Epithelial Stem Cells after Lung Injury. Stem Cell Reports 12, 1041–1055.

Zepp, J. A., Morley, M. P., Loebel, C., Kremp, M. M., Chaudhry, F. N., Basil, M. C., Leach, J. P., Liberti, D. C., Niethamer, T. K., Ying, Y., et al. (2021). Genomic, epigenomic, and biophysical cues controlling the emergence of the lung alveolus. Science 371.

Zepp, J. A., Zacharias, W. J., Frank, D. B., Cavanaugh, C. A., Zhou, S., Morley, M. P. and Morrisey, E. E. (2017). Distinct Mesenchymal Lineages and Niches Promote Epithelial Self-Renewal and Myofibrogenesis in the Lung. Cell 170, 1134–1148 e1110.

