## Supplemental figures for "Identification of a myofibroblast differentiation program during neonatal lung development"

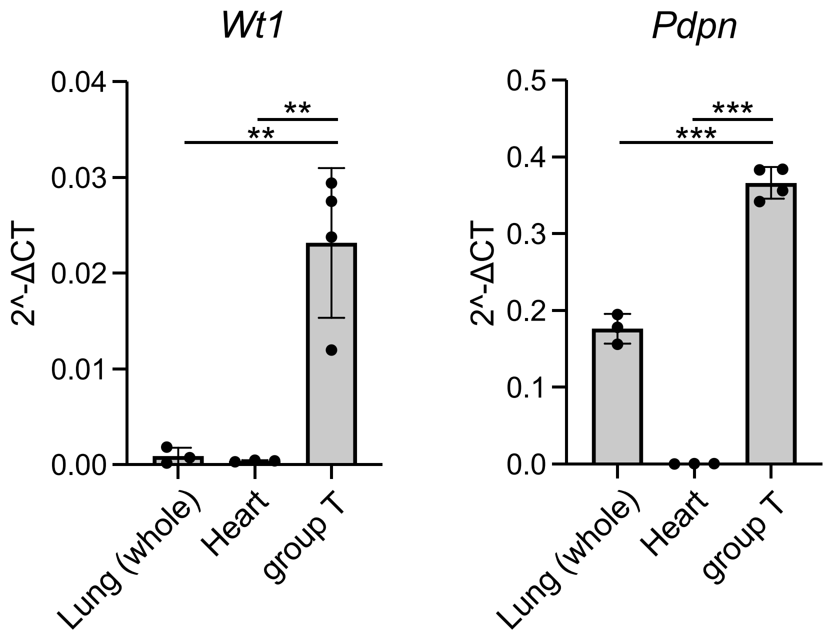


**Supplemental Figure 1. Quantitative RT-PCR of RNA from sorted T cells (TDTomato$^{+}$, GFP$^{-}$), whole lung, and whole heart.** T cells express the mesothelial marker Wilms tumor 1 homolog (Wt1) and the AT1 cell marker podoplanin (Pdpn, T1α). Wt1 was detected at low levels in whole lung and not detected in heart. Pdpn was expressed in whole lung but not in heart. n=3-4, ANOVA with Tukey's multiple comparison, ** $P$ < 0.002, *** $P$ < 0.0001.

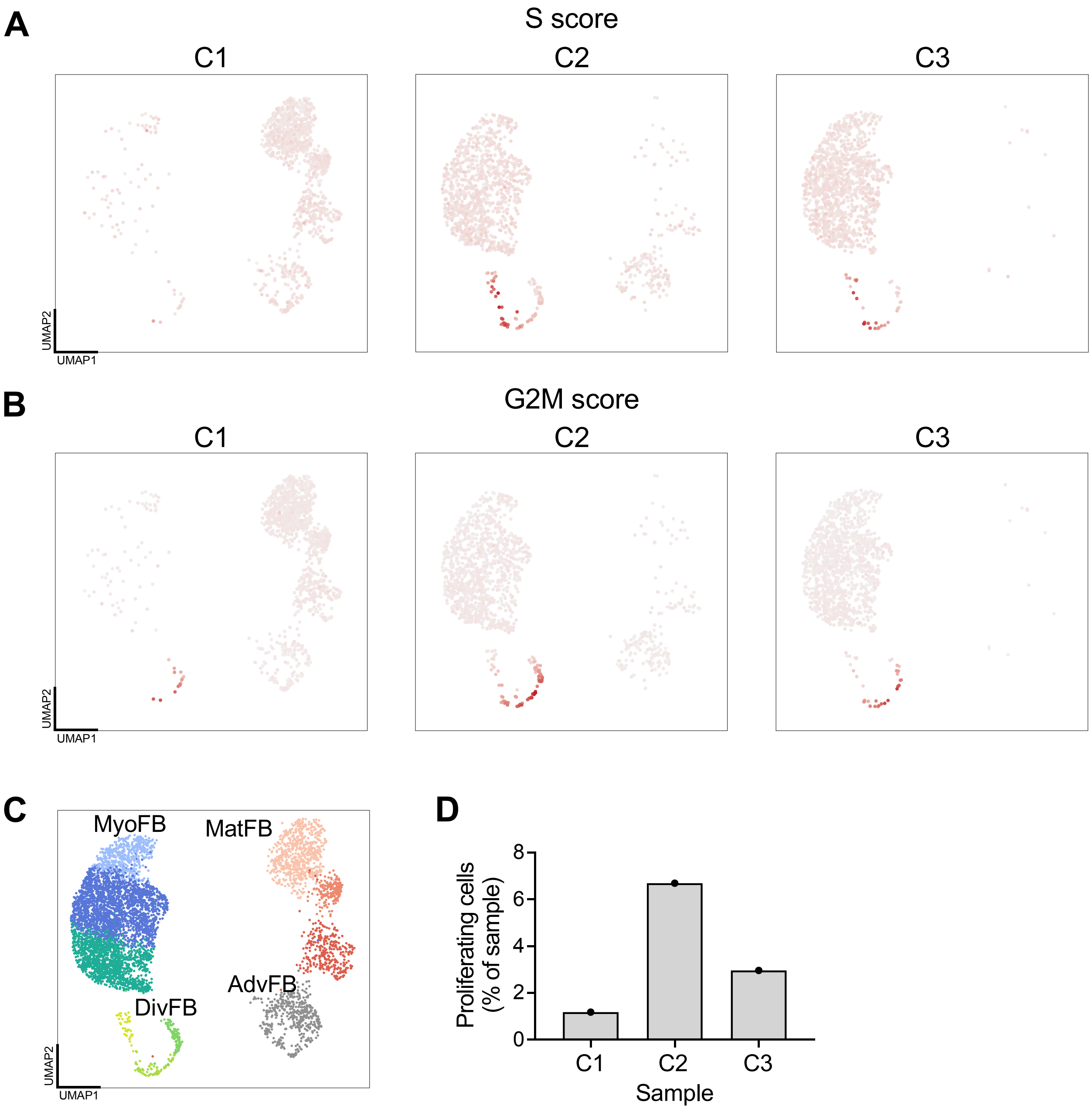


**Supplemental Figure 2. Analysis of cell cycle stages and distribution in sorted mesenchymal groups. A.** S phase cells, determined by Seurat Cell-Cycle Scoring are enriched in C2 cells and are present at low levels in C3 cells. **B**. G2/M phase cells are enriched in C2 cells and are present at low levels in C3 cells. **C**. reference UMAP showing primary cell types. **D**. Relative numbers of proliferating cells within primary sorted cell types normalized to the number of cells in each sample.

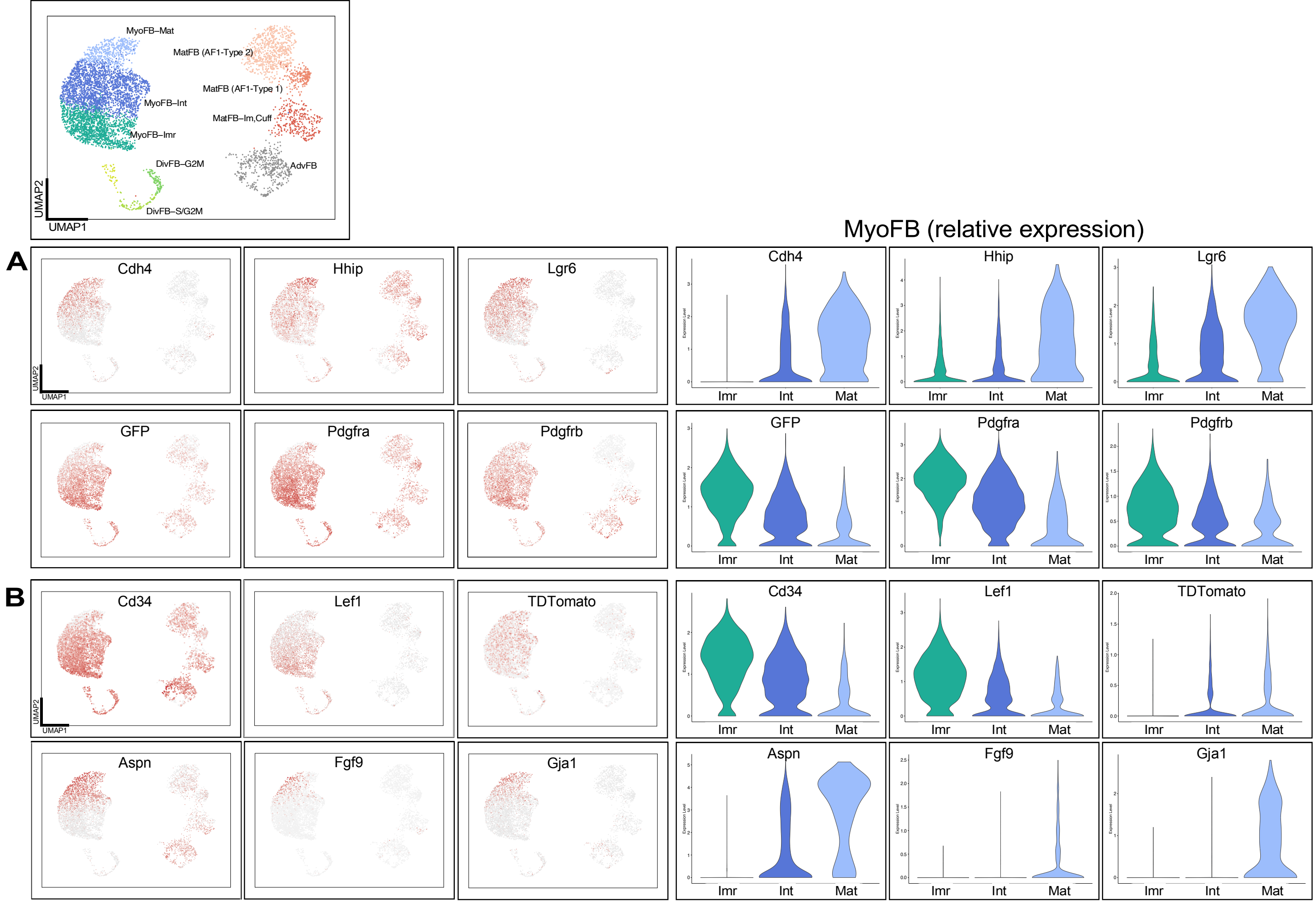


**Supplemental Figure 3. Identification of immature and mature myofibroblast subtypes. A**. UMAP and violin plots comparing myofibroblast subclusters in our data set with markers identified by Narvaez Del Pilar et al. that were present in both neonatal and adult lung (*Cdh4, Hhip*, and *Lgr6*) and *Pdgfra*-high expressing cells that were only present in neonatal lung. In our data set, *Cdh4, Hhip*, and *Lgr6* were associated with MyoFB-Mat cells and $Pdgfra^{EGFP}$ (GFP) and endogenous *Pdgfra-high* and *Pdgfrb* were associated with MyoFB-Imr cells. **B**. Genes associated with progenitor cells, *Cd34* and *Lef1*, were enriched in MyoFB-Imr cells whereas *Aspn, Fgf9* and *Gja1* were enriched in MyoFB-Mat cells. *Fgf18* lineage TDTomato was enriched in MyoFB-Int and MyoFB-Mat cells.

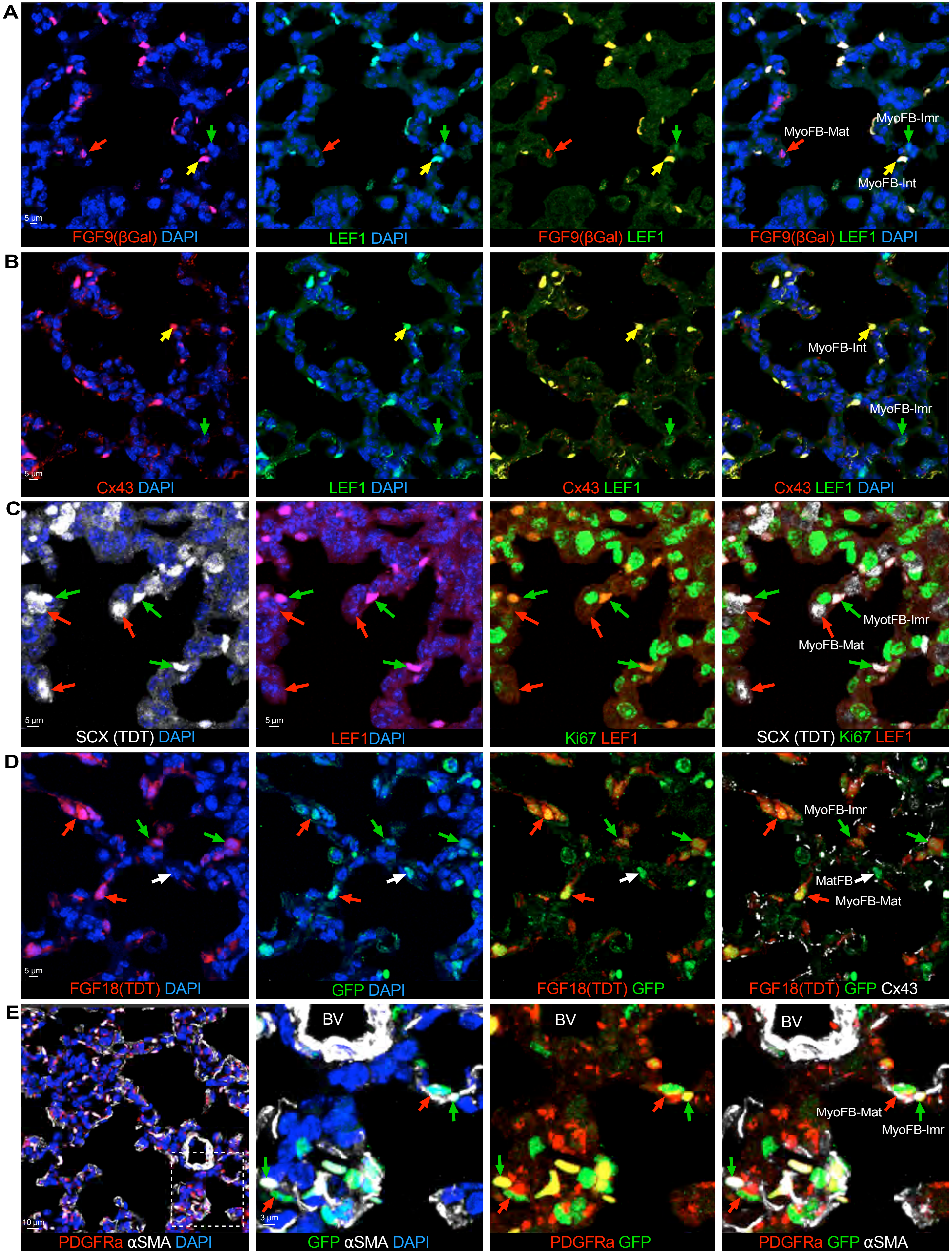


Supplemental Figure 4

**Supplemental Figure 4. Expression patterns of genes in the proximal and distal P7 mouse lung. A**. In the distal lung, FGF9 (βGal)$^{+}$, LEF1$^{-}$ (red arrow), FGF9 (βGal)$^{+}$, LEF1$^{+}$ (yellow arow), and FGF9 (βGal)$^{-}$, LEF1$^{+}$ (green arrow) cells were identified, corresponding to MyoFB-Mat, MyoFB-Int, and MyoFB-Imr cells, respectively. **B**. Cx43$^{-}$, LEF1$^{+}$ cells (green arrow) and Cx43$^{+}$, LEF1$^{+}$ cells (yellow arrow) correspond to MyoFB-Imr and MyoFB-Int cells, respectively. **C**. SCX (TDT)$^{+}$, LEF1$^{-}$, MKI67$^{-}$ cells (red arrows) and SCX (TDT)$^{+}$, LEF1$^{+}$, MKI67$^{+}$ cells (green arrows), correspond to MyoFB-Mat and MyoFB-Imr cells, respectively. **D**. FGF18 (TDT), $Pdgfra^{EGFP}$ (GFP)$^{High}$ cells correspond with MyoFB-Mat cells (red arrows); FGF18 (TDT)$^{Low}$, $Pdgfra^{EGFP}$ (GFP)$^{Low}$ cells correspond with MyoFB-Imr cells (green arrow); FGF18 (TDT)$^{-}$, $Pdgfra^{EGFP}$ (GFP), Cx43$^{+}$ cells correspond with MatFB (white arrow). **E**. Validation of endogenous PDGFRA and $Pdgfra^{EGFP}$ (GFP) expression. Dashed box in the left image indicates area shown at higher magnification in the following images. $Pdgfra^{EGFP}$ (GFP)$^{+}$ nuclear staining is localized in cells that have cytoplasmic staining for PDGFRA. $Pdgfra^{EGFP}$ (GFP)$^{Low}$, αSMA$^{+}$ cells are marked as MyoFB-Mat (red arrows); $Pdgfra^{EGFP}$ (GFP)$^{High}$, αSMA$^{+}$ cells correspond to MyoFB-Imr cells (green arrows).

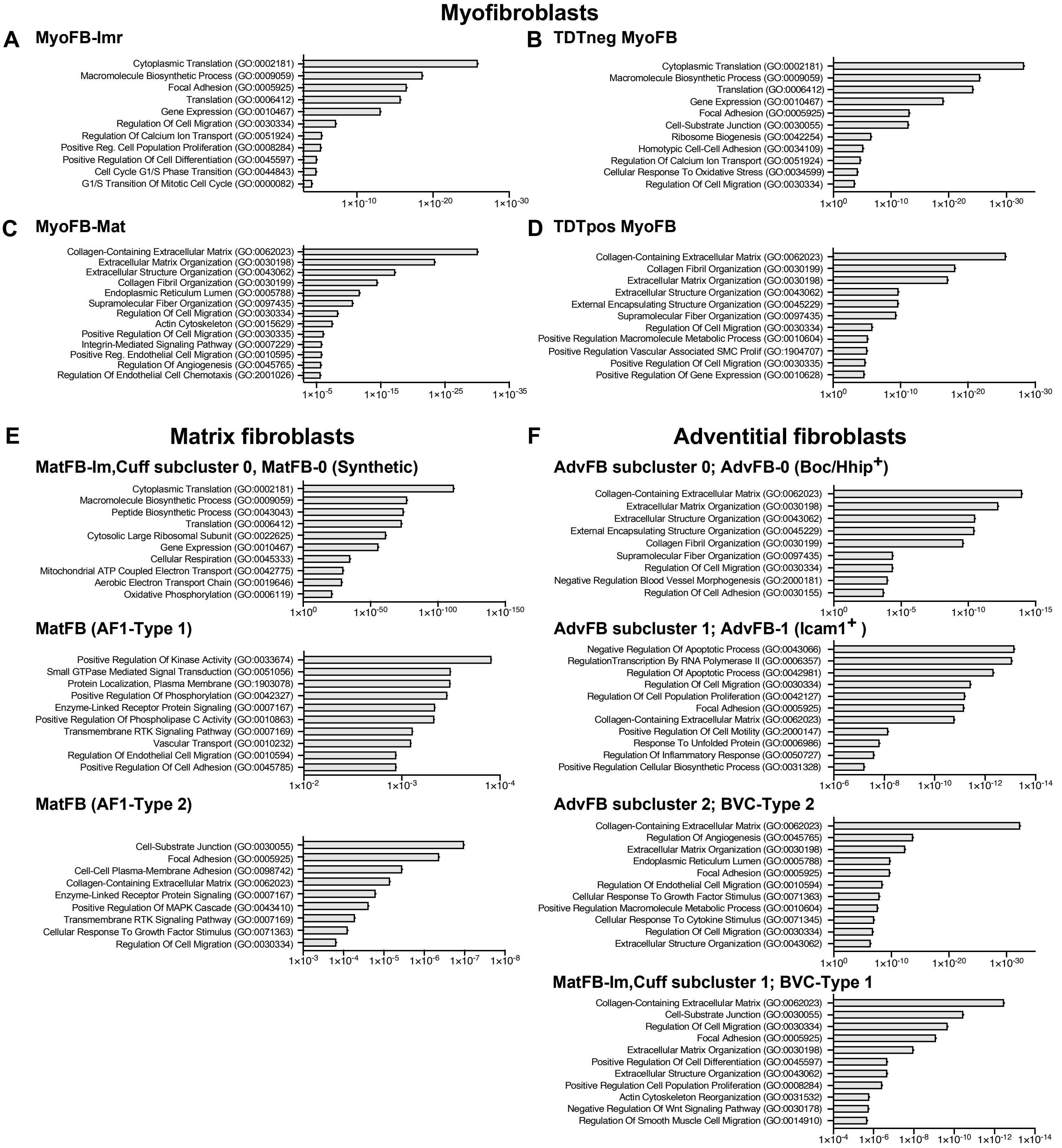


**Supplemental Figure 5. Gene Ontologies for differentially expressed genes in mesenchymal cell types.**

**A-D**. GO analysis of DEGs for MyoFB-Imr (**A**), MyoFB (TDTomato$^{-}$) (**B**), MyoFB-Mat (**C**), and MyoFB (TDTomato$^{+}$) (**D**). MyoFB (excluding proliferating cells) grouped by subcluster or grouped by TDTomato expression were used to identify DEGs using the FindAllMarkers function in Seurat (Log2FV > 0.25). **E**. MatFB (including subclusters Synthetic, AF1-Type 1, and AF1-Type 2) were used to identify DEGs using the FindAllMarkers function in Seurat (Log2FV > 0.25). **F**. AdvFB (including AdvFB subclusters 0, 1, 2 and BVC-Type 1) were used to identify DEGs using the FindAllMarkers function in Seurat (Log2FV > 0.25).

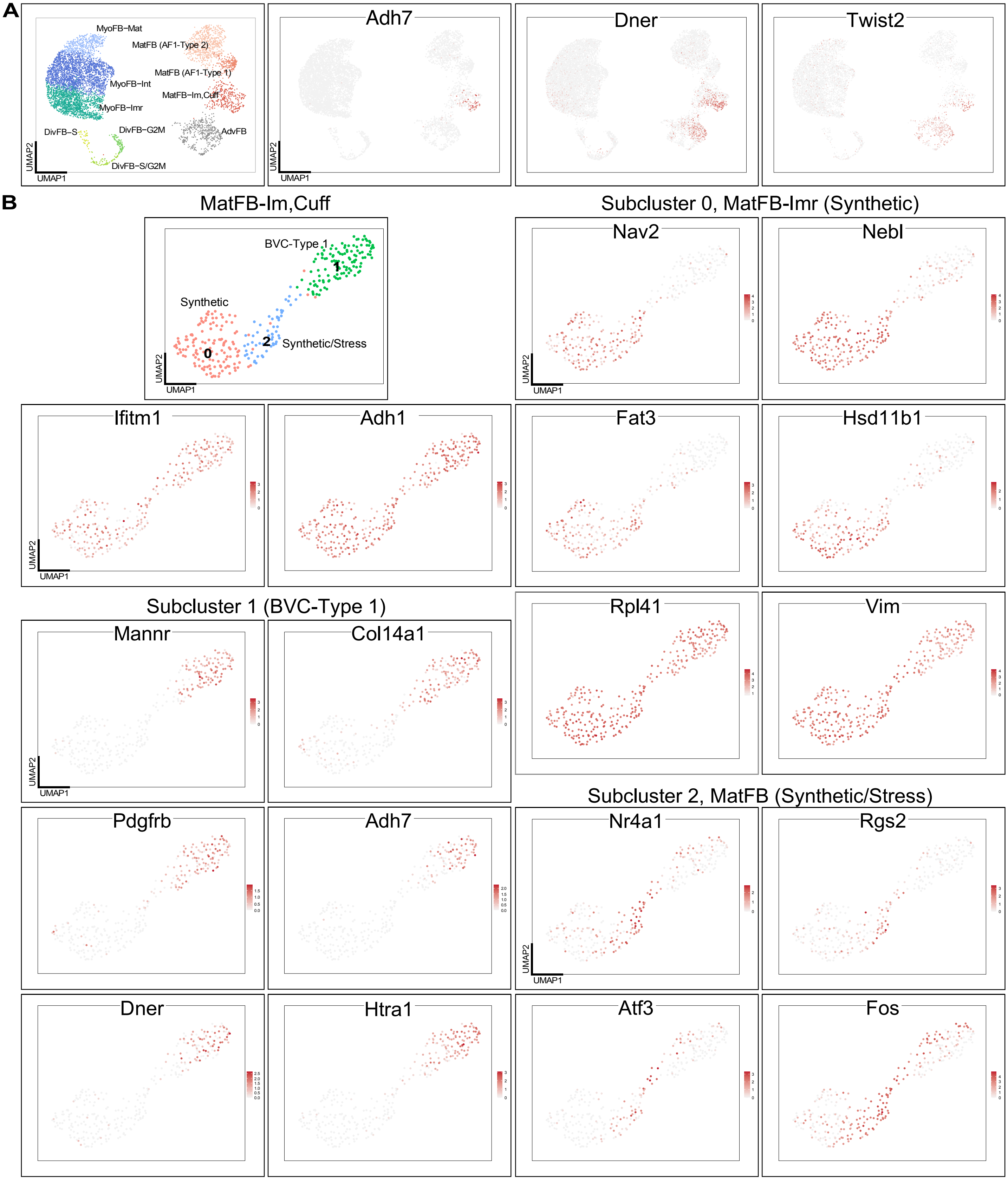


**Supplemental Figure 6. Subclustering of MatFB-Im,Cuff cells. A.** UMAP showing markers unique to MatFB-Im,Cuff cells and markers expressed in both MatFB-Im,Cuff and AdvFB cells. **B.** UMAP shows representative genes that are selectively expressed in subclusters of MatFB-Im,Cuff cells. Subcluster 0, designated MatFB-0 (synthetic), are enrichment for ribosomal and ECM genes and express high levels of *Nebl* and *Nav2*. Subcluster 1, designated BVC-Type 1, express *Adh7*, *Pdgfrb, Twist2, Dner* and *Col14a1* and corresponds to a subtype of an adult adventitial or bronchovascular cuff fibroblast. Subcluster 2 (Synthetic/Stress) expressed stress response factors such as *Fos*.

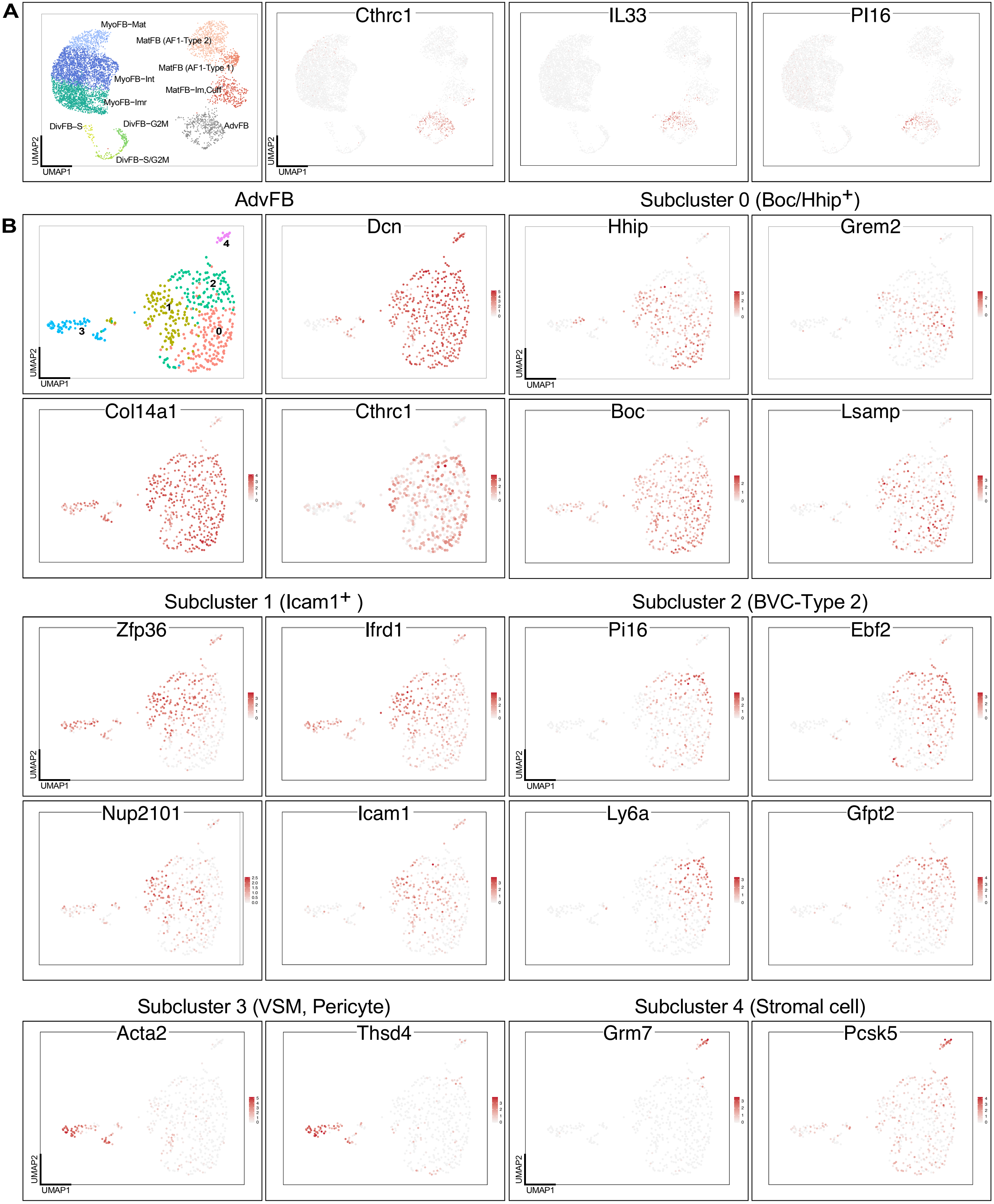


**Supplemental Figure 7. Subclustering of AdvFB cells. A.** UMAP showing markers unique to AdvFB cells. **B**. UMAP shows representative genes that are selectively expressed in subclusters. Subclusters 0 is enriched for expression of *Hhip, Boc,* and *Lsamp* and subcluster 1 is enriched for expression of *Zfp36* and *Icam1.* Subcluster 2, designated BVC-Type 2, express *Pi16* and *Ebf2* and corresponds to a subtype of adult lung adventitial fibroblasts. Subcluster 3 (VSM, pericyte) express *Acta2, Myh11, Tagln, Thsd4, Sost, Cbr2, Rgs5,* and *Cspg4* and is likely composed of vascular smooth muscle cells, a cell similar to vascular mural cells, and pericytes. Subcluster 4 expressed *Grm7*, *Pcks5, Musk,* and corresponds to an undefined lung stromal cell.

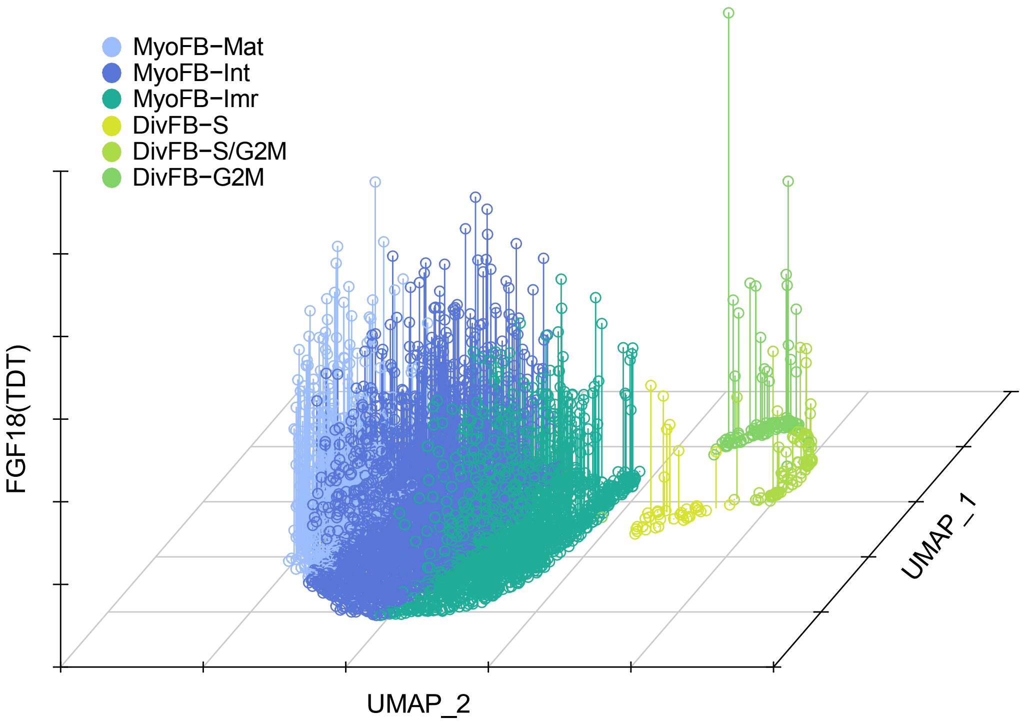


**Supplemental Figure 8. Expression level of TDTomato correlates with maturation of MyoFB**. UMAP of MyoFB clusters is plotted in the XY axis and FGF18 (TDT) expression is plotted in the Z axis.
